# Polyclonal and clonal organoid models of Barrett oesophagus and oesophageal adenocarcinoma reveal heterogeneity in progression and therapy response

**DOI:** 10.64898/2026.01.09.698383

**Authors:** Daniel H Jacobson, Dylan P McClurg, Yibing Wang, Emily Black, Charlotte Cassie, Tik Shing Cheung, Hannah Coles, Ginny Devonshire, SriGanesh Jammula, Benjamin A Hall, Xiaodun Li, Ahmad Miremadi, Krishnaa T Mahbubani, Massimiliano di Pietro, Kourosh Saeb-Parsy, OCCAMS Consortium, Rebecca C Fitzgerald, Christopher M Jones, Lizhe Zhuang

## Abstract

Oesophageal adenocarcinoma (OAC) is a major cause of morbidity and mortality. OAC and its precursor, Barrett oesophagus (BO), are defined by substantial early heterogeneity, complicating prevention and treatment of OAC. This diversity is poorly recapitulated by current *in vitro* and animal model systems. We have generated 91 patient- and healthy donor-derived organoids (PDOs) spanning normal oesophagogastric tissue (n=29), BO (n=13), dysplasia (n=8) and OAC (n=41). These PDOs capture population heterogeneity and recapitulate phenotypic, genomic and transcriptomic features of their respective disease stages. We develop a single cell-derived clonal organoid (sc-organoid) approach, with which we isolate 28 additional organoid lines representing subclonal populations that are difficult to discern and maintain in bulk PDO cultures, and which may represent clones at high risk of progression. Using this platform, we demonstrate functional utility across the pre-malignant to invasive disease spectrum, including recapitulation of step-wise disease progression, subclonal phylogenetic structure, and diverse responses of OAC to systemic anti-cancer therapies and radiotherapy.

**HIGHLIGHTS:**

1. Patient- and healthy donor-derived organoids (PDOs) provide a functional platform of disease progression and heterogeneity across normal gastric, non-dysplastic and dysplastic Barrett oesophagus (BO) and oesophageal adenocarcinoma (OAC).
2. We provide a quantitative phenotypic and molecular framework to assess the provenance and fidelity of each PDO model given the heterogeneity of this disease.
3. PDOs recapitulate key features of non-dysplastic and dysplastic BO, as well as invasive OAC.
4. scRNA-seq of PDOs across multiple disease stages reveals cellular heterogeneity and stage-specific cell clusters
5. Single cell-derived clonal organoids (sc-organoids) isolate and maintain high-risk subclonal populations.
6. OAC PDOs mirror known population level variation in response to systemic anti-cancer therapies.

## INTRODUCTION

Oesophageal adenocarcinoma (OAC) is an important cause of cancer-related morbidity and mortality, with an overall five-year survival rate of less than 20%^1^. Around 85,000 people are diagnosed with OAC each year, corresponding to a global age-standardised incidence of 0.9 cases per 100,000 persons^2^. Following increases in incidence since the 1970s, OAC is now the dominant subtype in over 20 high income countries, where rates reach as high as 3.4 cases per 100,000 people^2, 3^.

The development of OAC follows the replacement of the normal stratified oesophageal squamous epithelium with a metaplastic mosaic of gastric and intestinal like epithelium^4^. This lesion, Barrett oesophagus (BO), originates from cells within the gastric cardia, likely in response to exposure to injurious gastric refluxate^4, 5^. BO typified by intestinal metaplastic features can progress to dysplasia and invasive disease but this occurs in a minority of patients, with an annual risk of OAC of 0.3% for a person with a non-dysplastic lesion^6^. Greater phenotypic and molecular variation also correlates with the risk of progression to OAC^7–9^. A more precise understanding of events leading to progression is vital to enabling early detection and interception strategies^10^. The heterogeneous nature of the BO precursor lesion gives rise to a highly diverse cancer at phenotypic and molecular levels of resolution^11–13^. This tumour heterogeneity establishes a therapeutic landscape with few recurrent actionable drivers for targeted therapies and high levels of resistance to cytotoxic chemotherapy, external beam radiotherapy (EBRT) and immune checkpoint inhibition, leading to poor outcomes^10, 14^.

There is consequently a need to develop an experimental platform that recapitulates inter- and intra-lesional heterogeneity from the development of BO through to its dysplastic progression to OAC. In this context, it is challenging to model between-patient heterogeneity using animal models, for which there are also ethical and cost considerations ^15^. By contrast, we and others have previously shown that patient-derived organoids (PDOs) representing OAC can be established^16–19^; however, the extent to which these also reproducibly demonstrate heterogeneity in therapeutic response to established therapies including cytotoxic chemotherapy and EBRT is less certain^20, 21^. Moreover, successfully establishing PDOs that resemble precancerous lesions has proven more challenging^16^.

The aim of this study was to establish and characterise a PDO biobank of sufficient scale to recapitulate the full spectrum of heterogeneity at each stage of OAC carcinogenesis, including normal gastric epithelium, non-dysplastic and dysplastic BO, and treatment-naïve as well as treated OAC. Using a combination of genomic sequencing and *in silico* modelling of PDO growth and morphology, we provide a quantitative toolkit to characterise and phenotype each organoid. To address intralesional heterogeneity in BO, we also introduce novel, single cell-derived clonal organoids (sc-organoids) to unmask minor cell populations at high risk of dysplastic progression that may be missed in bulk analyses or lost through successive passages. Finally, we demonstrate that OAC PDO phenotypic and molecular heterogeneity translates to variation between PDOs in their response to different forms of cytotoxic chemotherapy, EBRT and targeted therapy.

## RESULTS

### Establishment of PDOs spanning BO and healthy comparison tissues

The first aim was to develop a robust and representative PDO resource spanning the entire spectrum of BO development and progression prior to OAC development. Thirteen PDOs were derived from non-dysplastic BO segments at routine surveillance endoscopy and eight PDOs were derived from punch biopsies from regions of dysplasia in endoscopic mucosal resection specimens **(Supplementary Table 1)**. Respective efficiencies for deriving PDOs were therefore 68% (13/18) and 61% (8/13) from non-dysplastic and dysplastic tissue. In addition,16 normal gastric PDOs were established, including seven from deceased healthy organ gastroesophageal squamocolumnar junction (n = 4) and gastric body (n=3) donor tissues **(Supplementary Table 2)**. Nine further gastric PDOs were derived from patients with BO: one each from the gastric body and antrum, and seven from the cardia to give a range of gastric lineages. Three further PDOs were derived from normal intestinal tissue as control for the intestinal phenotype, which characterises BO with malignant potential **(Supplementary Figure 1)**. All PDOs were passaged at least five times, and all were successfully revived following cryopreservation.

### BO PDOs recapitulate population heterogeneity and key features of non-dysplastic as well as dysplastic disease

We first sought to understand the genomic heterogeneity of BO PDOs in relation to dysplasia status and matched tissue samples. Median somatic mutation burden was comparable and reflected disease stage, with 2.11 muts/Mb in non-dysplastic PDOs versus 1.89 muts/Mb in matching tissues, and 5.76 muts/Mb in dysplastic PDOs versus 4.57 muts/Mb in matching tissues. Median burden in normal stomach was 0.11muts/Mb and 0.26 muts/Mb from healthy- and patient-derived PDOs respectively **(Supplementary Figure 2, A and B)**. As expected, PDOs showedhigher purity, defined as the proportion of epithelial cells in a sample, and higher clonality than their matching tissues, across non-dysplastic and dysplastic models **(Supplementary Figure 2, C and D)**.

A comprehensive overview of OAC driver genes, mutational signatures, and clinical annotations is provided in **Figure 1A**. As expected, there were few mutational events in normal gastric PDOs. In contrast, *CDKN2A* alterations, primarily deletions, were identified in 31% of non-dysplastic and 38% of dysplastic PDOs, consistent with the reported role of this gene in early BO development^22^. Alterations in *TP53* were seen in only one (1/13, 8%) non-dysplastic BO PDO but were more frequent (n=3/8; 37.5%) in PDOs taken from patients with dysplasia.

**Figure 1:**
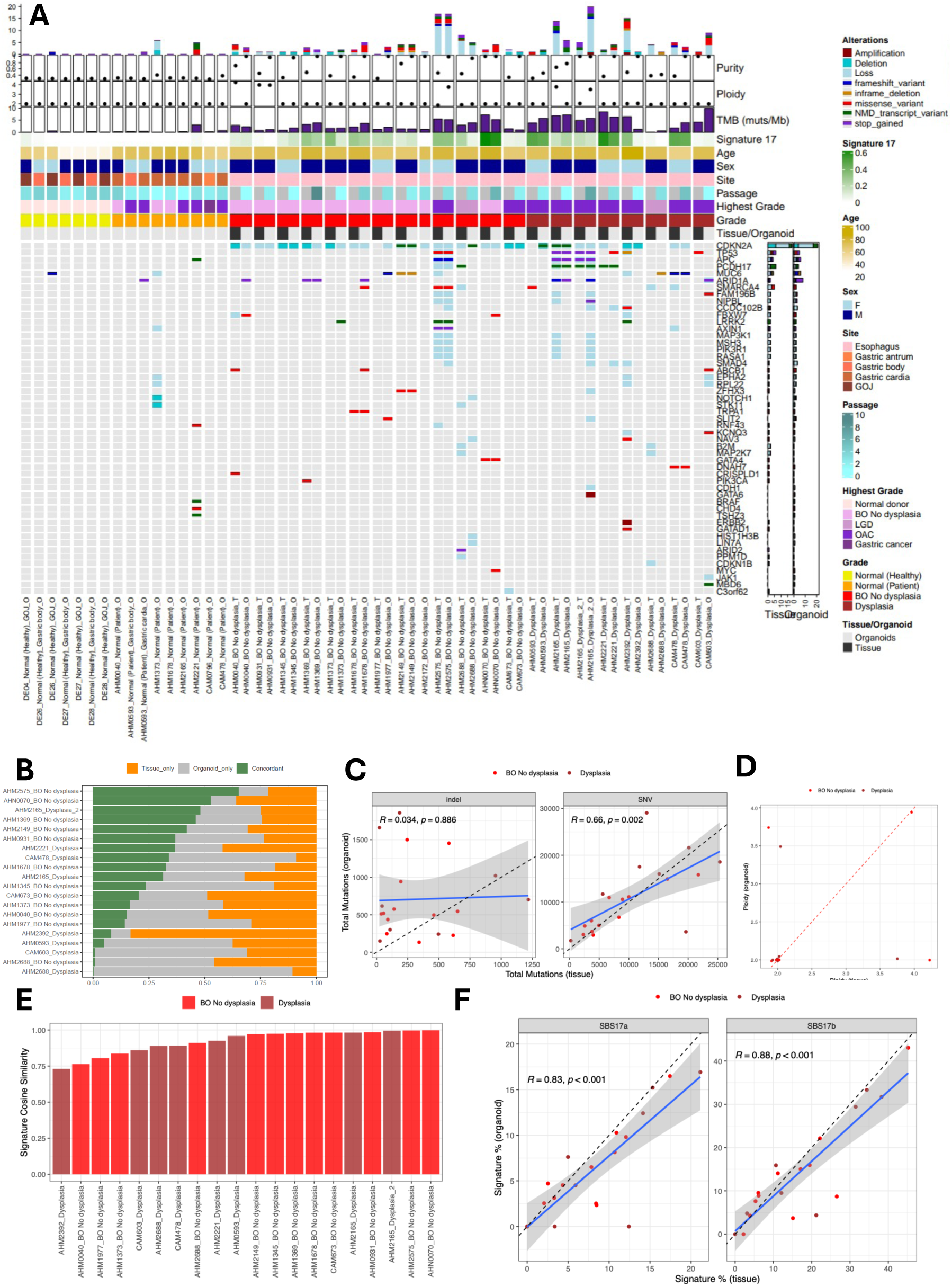
The genomic landscape of normal gastric and Barrett oesophagus-derived PDOs and its correlation with parent tissue samples. **(A)** Driver alterations and genomic features across PDOs and adjacent tissues. Driver genes were considered according to the set of genes (n=76) found to be positively selected in OAC by Frankell et al^11^ **(B)** Concordance of mutational profiles across Barrett oesophagus tissue-PDO pairs, defined by the proportion of mutations shared and unique across samples. Tissues from normal stomach did not undergo WGS. **(C-D)** Comparisons of **(C)** indel and SNV burdens, and **(D)** ploidy across tissue-PDO pairs. **(E)** Comparisons of mutational signature profiles across tissue-PDO pairs calculated by cosine similarity. **(F)** Correlations in SBS17a (left) and SBS17b (right) proportions across tissue-PDO pairs. Pearson correlation coefficients were used to determine correlation strengths.

The fraction of shared mutations between BO PDOs and their matching tissues ranged from 0.3-65.3% **(Figure 1B)**. Broadly, the mean fraction of private mutations (i.e. that are not shared) in PDOs was higher in dysplastic PDOs than in non-dysplastic PDOs (median 46.8% vs 38.1%). Whilst there was no correlation between the total number of insertion/deletion mutations in PDOs and matching tissue (r=0.034, p=0.89), there was a moderate correlation in single nucleotide variant (SNV) count (r=0.66, p=0.0015; **Figure 1C**) which was stronger for non-dysplastic PDO/tissue pairs (r=0.91, p<0.001) **(Supplementary Figure 2E)**. These results suggest increasing heterogeneity in dysplastic compared with non-dysplastic tissues which is thus reflected in the variation captured within a given PDO model.

Most tissues and PDOs were diploid; however, three PDOs captured whole genome doubling. This was mirrored in matching tissue for one PDO (AHM0931) but not in the other two (AHM2575 (non-dysplastic) and AHM2165 (dysplastic), which featured whole genome duplications despite diploid matching tissues **(Figure 1D)**. Overall concordance in homologous recombination deficiency (HRD) showed a moderate positive correlation for non-dysplastic pairs (r=0.52, p=0.081), but a dissimilar relationship for dysplastic pairs (r=–0.34, p=0.41) **(Supplementary Figure 2F)**. This suggests that it is challenging to capture chromosomal instability in PDOs and one cannot rely on the genomic profile of matching tissues.

In comparison to the accumulation of driver gene mutations, mutational signatures are laid down early in the pathogenesis of BO and maintained consistently through malignant progression^24, 25^. In keeping with this, we observed a high concordance of mutational signatures in the PDOs and matching tissues, with a median cosine similarity of 0.97 (range 0.73-1.00), which did not differ between dysplastic and non-dysplastic PDO-tissue pairs **(Figure 1E and Supplementary Figure 2, G and H).** In particular, single base substitution signature 17 (SBS17a/b), a characteristic feature of BO and OAC, showed a strong overall correlation for both SBS17a (r=0.83, p<0.001) and SBS17b (r=0.88, p<0.001) **(Figure 1F, Supplementary Figure 2I)**. As expected, SBS17a/b proportions were similar across non-dysplastic and dysplastic PDOs **(Supplementary Figure 2J)**.

We also observed clonal heterogeneity and drift in precancerous PDOs **(Supplementary Figure 2, K to M).** For example, AHM0040 contained a major subclonal population (VAF 18%) harbouring *ARID1A* and *FBXW7* driver mutations. In AHM1373, a subclonal cluster (median VAF ∼19%) was present at passage 1 but became clonal by passage 6 (median VAF 50%). Whilst no driver alterations were present in both passages, passage 1 harboured a mutation in *LRKK2* **(Figure 1A**), whilst passage 6 carried frameshift variants in *ARID1A*, *ARID1B*, and *MUCc*, possibly indicating selection of a separate, rarer subclone. Hence, these models evolve temporally, and this can be tracked genetically as required for their utility to study disease progression and interception.

### Single cell-derived BO PDOs recapitulate intralesional heterogeneity

In view of the substantial genomic heterogeneity and the passage-related clonal drift, we reasoned that isolating low-frequency, high-risk populations would enable single clone modelling of BO progression. To achieve this, we developed a microdissection method to derive individual organoids from bulk PDO culture. We termed these single cell-derived clonal organoids (sc-organoids) **(Figure 2A).**

**Figure 2:**
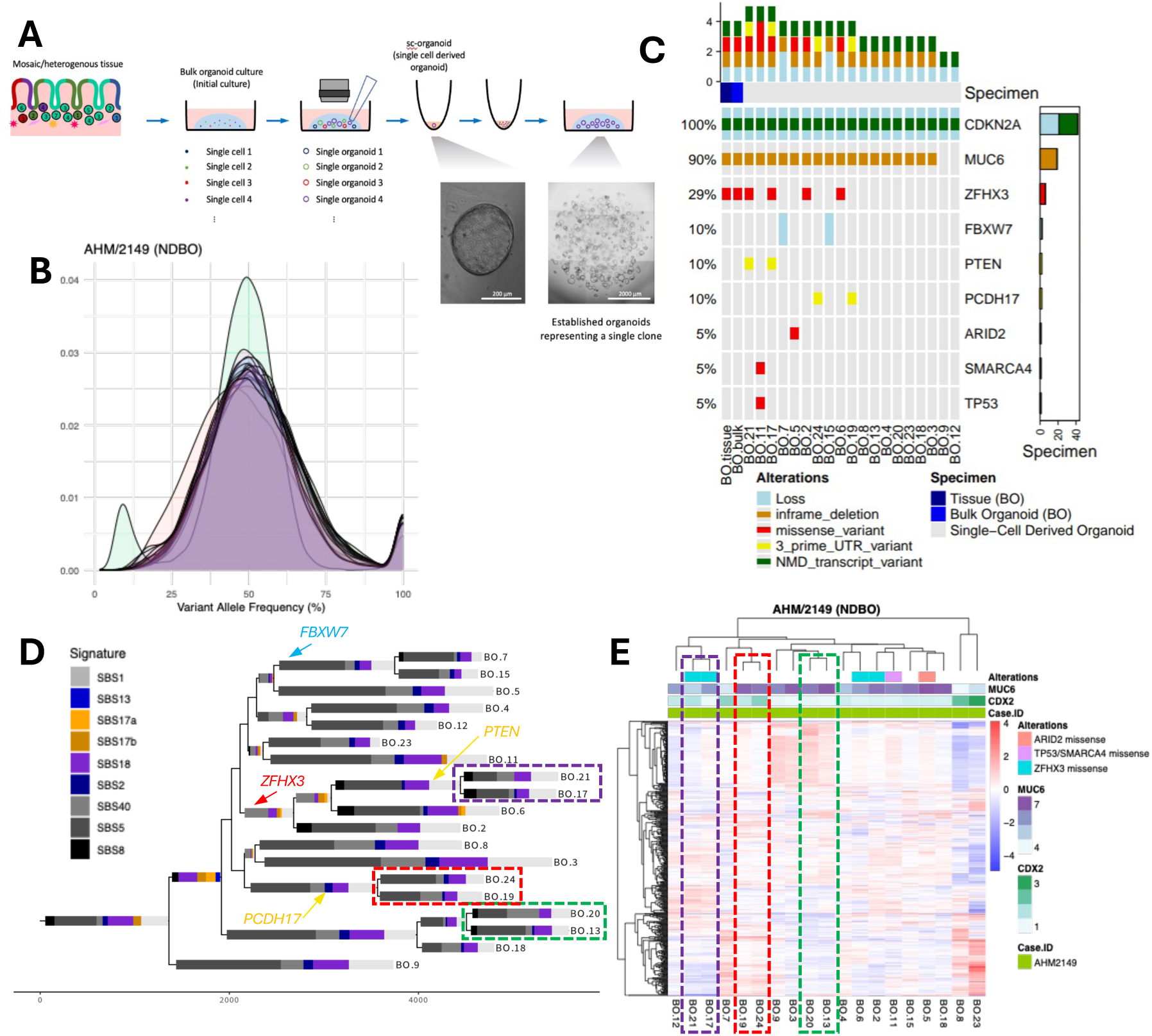
Diversification of non-dysplastic Barrett oesophagus demonstrated in a single cell-derived clonal organoid panel. **(A)** Approach for establishing sc-organoids. **(B)** Distributions of VAFs from single cell-derived organoids (n=19) derived from AHM2149. **(C)** Driver alterations found across the single cell-derived clonal organoid panel and adjacent bulk PDO and tissue. **(D)** A phylogenetic profile of organoids derived from AHM2149 defined using the sc-organoid panel. Signature contributions were calculated across mutations contributing to each branch respectively. Branch lengths are generally proportional to total amount of mutations. Signatures were not fitted for branches with fewer than 25 SNVs. **(E)** Hierarchical clustering of RNA-seq profiling for the sc-organoid panel using the top 500 variable genes following variance stabilised transformation of read counts. Heatmap annotation includes genetic alterations as reported in **(C)** and log2-transformed TPM-normalised counts of *MUCc* (a gastric marker) and *CDX2* (an intestinal markers), the correlation of which is reported in Supplementary Figure 6E. The coloured dashed squares found in **(D)** and **(E)** correspond to adjacent clones found through phylogenetic analysis and unsupervised clustering of RNA-seq.

We first generated 19 sc-organoids from a non-dysplastic case (AHM2149) and confirmed their clonality by analysing VAF distributions (a narrow peak at 50% indicates clonality) **(Figure 2B)**. The sc-organoid panel recapitulated key genomic features of matching BO tissue as well as the related bulk PDOs, including deletions of *CDKN2A* and *FBXW7,* and mutations in *MUCc* and *ZFHX3* **(Figure 2C)**. All sc-organoids contain private mutations (3.44%-45.8%), confirming that each represents a distinct lineage with preserved mutational signatures **(Supplementary Figure 3, A and B)**. Interestingly, one sc-organoid, BO.11, harboured mutations in *TP53* and *SMARCA4* that were not observed through WGS of the bulk PDO nor its matching tissue, suggesting a potential “seed” for dysplastic progression.

Since the sc-organoids were microscopically isolated from bulk PDO cultures only six days after tissue dissociation and seeding (see Establishment of single cell-derived clonal organoids, **Methods**), the clonal driver mutations identified in the sc-organoids are extremely unlikely to be acquired during culture and instead represent pre-existing small clones within the original tissue. Supporting this, we used deep targeted amplicon sequencing to confirm the presence of the *TP53* mutation in the bulk PDO, together with other driver mutations, including *SMARCA4*and *PCDH17*, that were not detected by WGS (**Supplementary Figure 3C**). Other mutations in *ARID2* and *PCDH17*, previously identified within individual sc-organoids (BO.5, BO.19 and BO.24), were also found in either the bulk PDO or the matching tissue, overall demonstrating that the sc-organoid panel is able to model the heterogeneity in BO that may not be adequately captured by WGS. It is noteworthy that not all mutations are detected in the matching tissue, though this is perhaps unsurprising given that heterogeneity and clonal expansion in BO occur over millimetre scales **(Supplementary Figure 3E)**^26^.

We next assessed the relatedness of the sc-organoids by constructing a phylogenetic tree using WGS data **(Figure 2D)**. Most mutations (85.98%) fitted the tree perfectly with minimal homoplasy, and driver alterations showed good concordance within the branching structure, reflecting their individual acquisition along the evolutionary trajectory. For example, *FBXW7* loss was shared by BO.7 and BO.15 but absent in the bulk PDO, whereas a *ZFHX3* mutation was detected in the bulk PDO but found in only four closely related sc-organoids.

To determine whether closely related sc-organoids also display phenotypic similarity, we compared transcriptomic concordance with WGS-inferred phylogenies (**Figure 2E**). We also assessed whether sc-organoids represent distinct lineages by analysing enrichment of individual lineage markers (*MUCc* versus *CDX2*) and lineage gene panels. Across 19 non-dysplastic BO sc-organoids, *MUCc* (gastric) and *CDX2* (intestinal) expression were strongly negatively correlated (r = -0.71; p = 0.00075, **Supplementary Figure 3D**), indicating that individual sc-organoids tend to adopt either a gastric or intestinal state. Notably, two sc-organoids (BO.8 and BO.23) displayed diverging transcriptional profiles to the rest of the sc-organoids, with significantly higher *CDX2* expression (**Figure 2E**).

With these encouraging data we generated a panel of nine sc-organoids from dysplasia tissue from an early cancer case (AHM2392), from which a bulk PDO shared only 8.15% of SNVs and indels **(Figure 1B),** to better understand and model clonal heterogeneity. All sc-organoids appeared clonal, except for one (BO.14) that also harboured a nonsense mutation in *APC* present at a VAF of 25% **(Supplementary Figure 4A),** suggesting it may be a mixture of two clones. Taken together these sc-organoids shared a large proportion of mutations with the matching bulk PDO **(Supplementary Figure 4B)**, whilst also presenting with more mutations than the bulk PDO on account of their high clonality **(Supplementary Figure 4C)**. The individual sc-organoids nevertheless demonstrated clonal diversity **(Supplementary Figure 4D)** and transcriptomic heterogeneity **(Supplementary Figure 4E)**, which with the isolation of an *APC*-mutant clone reaffirms the potential for this technique to isolate high-risk populations not captured in bulk PDOs **(Supplementary Figure 4F)**.

### Time-lapse imaging and *in silico* modelling illustrate morphological and growth kinetic differences in BO PDOs

The genomic profile must be interpreted in the context of PDO morphology and growth patterns to understand how this translates to phenotype. To assess morphology, we genetically engineered non-dysplastic (AHM0040) and dysplastic (AHM2221) PDOs with nuclear fluorescent tags (H2B-GFP/mCherry) and imaged using time-lapse microscopy. Unlike the characteristic solid morphology of OAC PDOs^16^, non-dysplastic PDOs showed four characteristic morphologies: a single cell layer surrounding a single lumen, a single cell layer with multiple lumens, multiple cell layers with a lumen and a solid morphology with no visible lumen **(Figure 3, A to C)**. Luminal PDOs expanded over time, whereas solid structures showed minimal growth **(Supplementary Table 3).** In contrast, dysplastic PDOs lack single cell layer multiple lumen morphology and have a higher proportion of solid morphology compared with non-dysplastic PDOs **(Figure 3D, Supplementary Figure 5, A to C; Supplementary Table 4)**.

**Figure 3:**
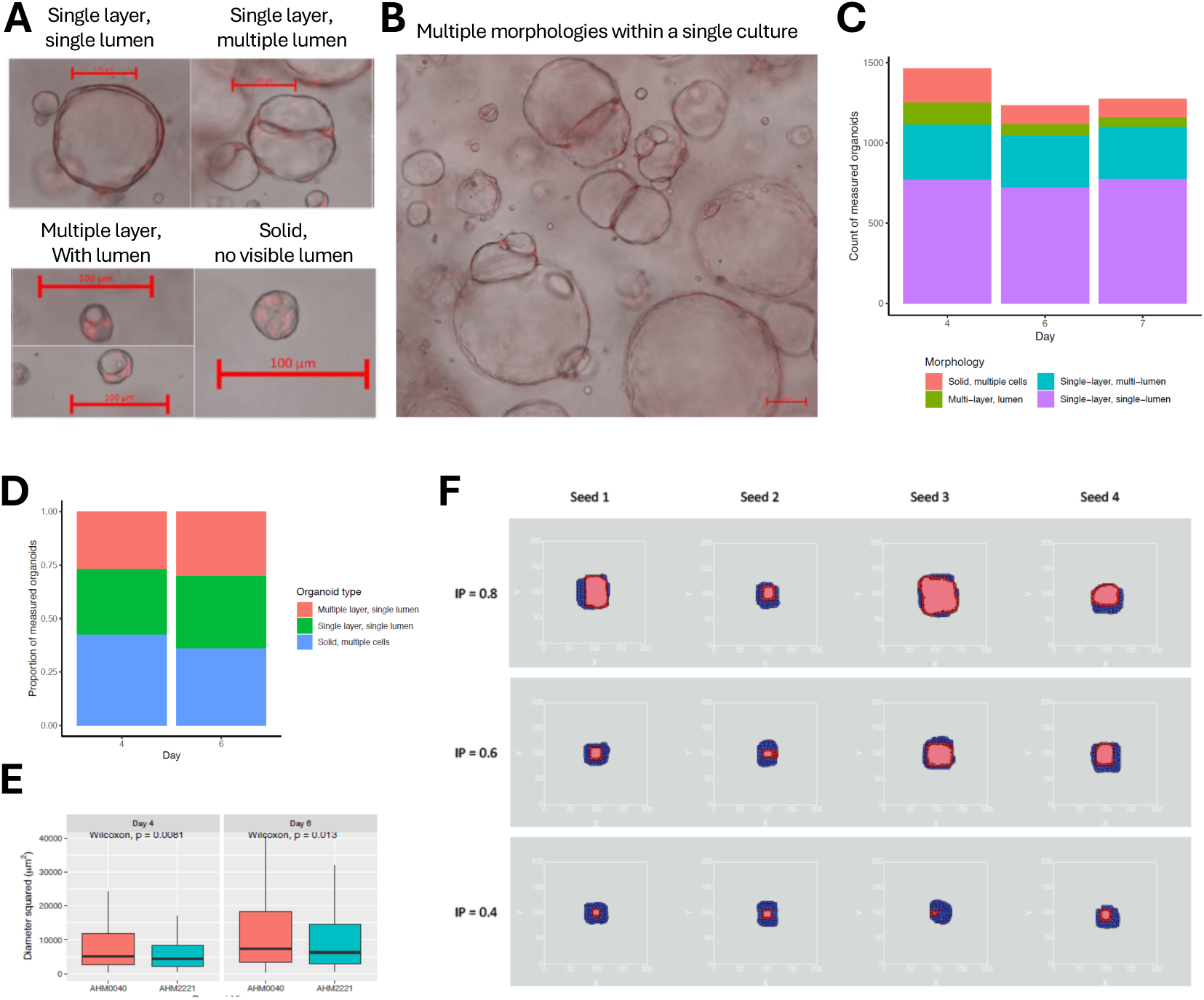
Analysis of key morphological features observed in Barrett oesophagus PDOs. **(A)** Representative bright-field images showing the four morphology classes: a single cell layer surrounding a single lumen, a single cell layer with multiple lumens, multiple cell layers with a lumen and a solid morphology with no visible lumen. In all instances the red scale bar represents a length of 100µm. **(B)** Example field of view illustrating the coexistence of multiple morphologies within a single dome of non-dysplastic (AHM0040) Barrett oesophagus PDOs. Scale bar represents 200µm. **(C)** Stacked bar plot of the count of each morphology class over seven days of growth following seeding in non-dysplastic (AHM0040) PDOs. **(D)** Proportional composition of morphologies at the indicated time points (normalised to total organoids measured per day) for dysplastic (AHM2221) PDOs. **(E)** Diameter squared (proportional to number of cells) of non-dysplastic (AHM0040) and dysplastic (AHM2221) PDOs at days 4 and 6 following seeding. **(F)** *In silico* simulations of organoid growth/morphology across four random seeds under three scenarios depicting variable likelihood of inheriting polarity (IP).

We observed minimal growth in almost all solid non-dysplastic organoids but visible expansion in luminal types, measured by both organoid size and nuclear number **(Supplementary Figure 5, D and E)**. Moreover, multiple and single lumen non-dysplastic PDOs demonstrated a similar, linear growth rate with considerable variation in individual PDO growth **(Supplementary Figure 5, F and G)**. Growth kinetics of single-lumen PDOs were comparable between non-dysplastic and dysplastic models **(Figure E, Supplementary Figure 5H)**. Time-lapse imaging also suggested that multi-lumen PDOs might be a result of fusion events from single luminal PDOs, providing a novel insight into the changing clonality of bulk organoid culture over time **(Supplementary Figure 5, I and J).**

Lumen formation has been linked to polarity, the loss of which is associated with dysplastic progression^27^. We established a lattice-based multi-site Cellular Potts Monte Carlo *in silico* model of organoid growth in which organoid polarity, and the probability of a daughter organoid exhibiting polarity, were varied **(Supplementary Figure 6A)**. Using this model, we demonstrate that a single layer single lumen morphology is acquired where organoids and their progeny demonstrate polarity, whereas organoids without polarity develop solid non-luminal structures **(Supplementary Figure 6B)**. Variation in the probability of acquiring polarity results in a variability in lumen formation **(Figure 3F)**. The model also accurately recreated the effects of observed fusion processes **(Supplementary Figure 6, C to E)** and captured the loss of polarity in dysplastic and OAC PDOs **(Supplementary Table 5).**

### Barrett oesophagus PDOs offer a functional platform for premalignant studies

To extend the phenotypic characterisation we used transcriptomic profiling based on lineage markers determined from our previous single cell RNA sequencing (scRNA-seq) datasets^5, 28^. As expected, the majority of BO PDOs demonstrated an intermediate phenotype between those seen for gastric and intestinal PDOs **(Figure 4A, Supplementary Figure 7, A and B)**. When comparing the bulk transcriptomes of normal and precancerous PDOs using a principal component analysis (PCA), BO and intestinal PDOs show a clear separation from gastric PDOs, in keeping with the acquisition of an intestinal phenotype **(Figure 4B)**. When comparing differentially expressed genes (DEGs) between three paired comparisons: gastric versus non-dysplastic PDOs, gastric versus dysplastic PDOs, and non-dysplastic versus dysplastic PDOs, broad concordance was observed in non-dysplastic and dysplastic PDOs when compared with gastric, while a small number of genes distinguished dysplastic and non-dysplastic PDOs **(Figure 4, C and D, Supplementary Figure 7, C to E)**. Pathway analyses revealed that upregulated genes in both non-dysplastic and dysplastic PDOs were enriched for developmental, migratory, proliferative, and intestinal differentiation programs **(Figure 4E, Supplementary Table 6)**, whereas down-regulated genes mapped to cell-peripheral processes and transcription factor binding sites **(Figure 4E, Supplementary Table 6)**. In addition, upregulation of extracellular matrix–related genes were specifically enriched in non-dysplastic PDOs **(Supplementary Figure 7F, Supplementary Table 7),** while in dysplastic PDOs, down-regulated genes mirrored the transcription factor binding site enrichment observed in the shared down-regulated profiles **(Supplementary Figure 7G, Supplementary Table 8)**. Intriguingly, three *HOX* genes (*HOXB2, HOXBc, HOXB8*), showed increased expression in dysplastic versus non-dysplastic PDOs **(Figure 4, D and F)**, with a similar pattern also observed for *HOXB5* and *HOXB7* **(Supplementary Figure 7H)**. This is consistent with our prior work demonstrating a role for HOX genes in OAC carcinogenesis^29^.

**Figure 4:**
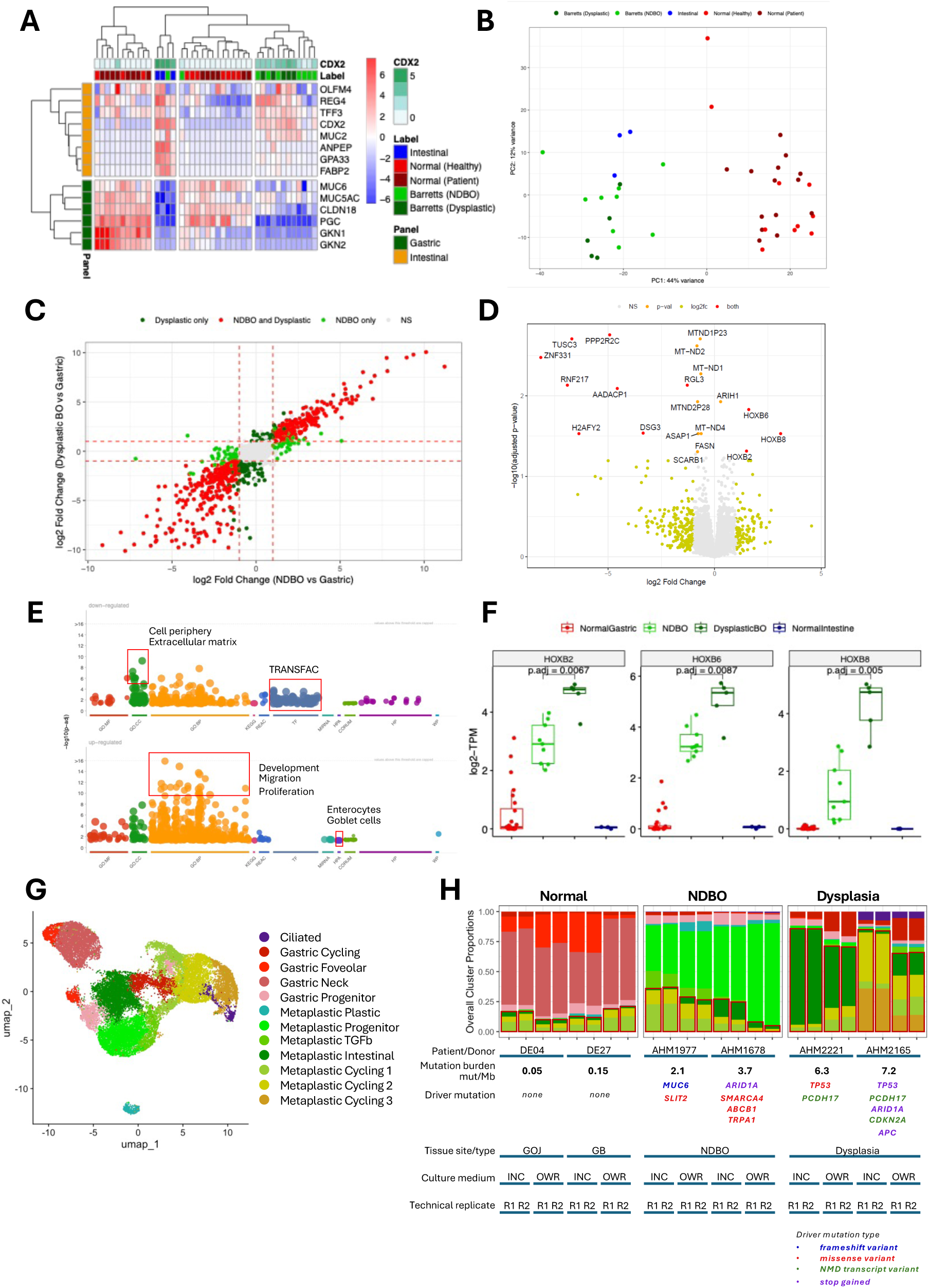
Transcriptomic profiling of control normal gastric and intestinal, and Barrett oesophagus, PDOs. **(A)** Inter-PDO expression of gastric- and intestinal-related genes. The reported scores were calculated using Z-normalisation following log2-transformation of TPM values. **(B)** Principal component analysis of all normal- and Barrett oesophagus PDOs based on the top 500 variable genes following variance stabilising transformation. **(C)** Correlations of log2-fold changes from differential expression analysis on non-dysplastic Barrett versus normal gastric, and dysplastic Barrett versus normal gastric PDOs. Genes were considered significant if they had an adjusted p-value < 0.05 and absolute log2-fold change > 1, as indicated by the dashed red lines. Full results of these analyses are reported in **Supplementary Figure 5D-E**. **(D)** Differential expression across dysplastic (n=5) versus non-dysplastic (n=9) Barrett oesophagus PDOs. Positive log2-fold change refers to increased expression in the dysplastic PDOs. Genes were coloured based on their adjusted p-value being less than 0.05, and absolute log2-fold change greater than 1. **(E)** Functional enrichment analysis of up- and down-regulated genes shared across non-dysplastic and dysplastic PDOs compared with normal gastric PDOs. Full functional enrichment analysis results are reported in **Supplementary Table 6**. **(F)** Expression of *HOXB2*, *HOXBc*, and *HOXB8* across all control normal and Barrett oesophagus PDOs. Values reported are log2-transformed TPM values, and comparison was done on the non-dysplastic and dysplastic PDOs using a two-sided Wilcoxon test with Benjamini-Hochberg adjustment for multiple testing. **(G)** UMAP dimensionality reduction summary of scRNA-seq analysis of 24 PDOs derived from normal gastric, BO, and dysplasia samples, labelled by assigned cell phenotypes. **(H)** Breakdown of present cell populations across 24 PDOs analysed by scRNA-seq. Metaplastic Intestinal, Metaplastic Cycling 1, 2 and 3 are grouped and labelled by red boxes.

### Single cell RNA sequencing of PDOs reveals stage specific cellular heterogeneity

To further dissect the phenotypic heterogeneity of PDOs, scRNA-seq was conducted on six PDO lines spanning three disease stages: (i) normal gastric: DE04 (0.05 mut/Mb) and DE27 (0.15 mut/Mb); (ii) non-dysplastic BO: AHM1977 (2.1 mut/Mb*, MUCc* frameshift, *SLIT2* missense), AHM1678 (3.7 mut/Mb*, ARID1A* stop-gained, *SMARCA4* missense, *ABCB1* missense, *TRPA1* missense); and (iii) dysplasia: AHM2221 (6.3 mut/Mb*, TP53* missense, *PCDH17* NMD transcript variants), AHM2165 (7.2 mut/Mb*, TP53* stop-gained, *ARID1A* stop-gained, *CDKN2A* NMD transcript, *APC* stop-gained, *PCDH17* NMD transcript variants) (see **Figure 1A** and **4H**). PDOs were cultured in technical duplicate under two media conditions, IntestiCult (INC), a commercially available medium used to establish PDOs from tissues, and OWR, the classic WNT- and R-spond in-based medium used to maintain PDOs (see **Methods**). Following alignment, processing, and quality control, a total of 26,583 cells were analysed (**Figure 4G, Supplementary Figure 8, A and B**).

Twelve clusters were identified, broadly falling within mosaic gastric and intestinal lineages consistent with the known cellular composition of BO (**Supplementary Figure 8C**). Clusters with clear lineage markers included Metaplastic Intestinal (*REG4*, *TFF3*, *CDX2*, *GPA33*), Gastric Neck (*MUCc*), and Gastric Foveolar (*CLDN18*, *GKN1*, *GKN2*). For clusters with less distinct lineage identities, *HOXB* gene expression was used as an additional discriminator, supported by our bulk RNA-seq data (**Figure 4F, Supplementary Figure 7H**) and consistent with published literature: negative *HOXB* expression indicates gastric lineage, while positive expression indicates metaplastic lineage^29^. Using this approach, we defined Gastric Progenitor (*CD44*, *HOXB*-negative), Metaplastic Progenitor (*CD44*, *HOXB*-positive), Metaplastic TGFβ (*TGFB1*, *HOXB*-positive), and EMT-like Metaplastic (*SNAI1*, *DLX2*, *HOXB*-positive) clusters. Four cycling clusters were identified based on proliferation and cell-cycle markers (*MKIc7, TOP2A, CENPF, CDC20*), Metaplastic Cycling 1, 2 and 3, and Gastric Cycling, corroborated by cell-cycle phase profiling (**Supplementary Figure 8D**). A Ciliated cluster (*ZMYND10*) was also identified, consistent with a cluster previously reported in a Barrett’s scRNA-seq dataset^30^.

Normal gastric PDOs were almost exclusively composed of gastric clusters, including Gastric Neck and Foveolar (**Figure 4H**). Notably, a small proportion of metaplastic clusters was also present, which we attribute to the plasticity of gastric cells in organoid culture and is consistent with our broader conclusion that metaplastic cells originate from a gastric progenitor^5^. BO and dysplastic PDOs were dominated by metaplastic populations. The Metaplastic Progenitor cluster was preferentially associated with non-dysplastic BO PDOs, while Metaplastic Intestinal and Metaplastic Cycling clusters were significantly enriched in dysplastic PDOs. Interestingly, Metaplastic Intestinal was predominantly associated with AHM2221, whereas Metaplastic Cycling was predominantly associated with AHM2165. We reason this may reflect different substages of dysplasia: though both PDO lines were derived from histopathologically confirmed HGD tissue, AHM2221 represents HGD alone, while AHM2165 was derived from HGD tissue adjacent to OAC, and harbours a higher mutation burden and greater number of driver mutations.

Notably, scRNA-seq findings were consistent with bulk RNA-seq data for *HOXB* genes; *HOXB8* in particular was markedly elevated in dysplastic PDOs relative to non-dysplastic BO (**Supplementary Figure 8G**). Both INC and OWR media gave rise to broadly similar cellular compositions, though OWR appeared to preferentially support Gastric Cycling and Neck cell populations, while INC tended to support Metaplastic Cycling populations (**Supplementary Figure 8, E and F**). Collectively, these scRNA-seq analyses demonstrate that PDOs from different disease stages retain distinct cellular compositions, with enrichment of metaplastic intestinal and cycling populations as a feature of dysplastic PDOs.

### OAC PDOs recapitulate population-level molecular heterogeneity with variable growth behaviours

The phenotypic and molecular heterogeneity that defines Barrett oesophagus is also a characteristic feature of OAC^13^. Expanding on our previous small panel of 10 OAC PDOs^16^, we show here that a biobank of 41 PDOs (donor characteristics outlined in **Supplementary Table 11**), recapitulates the genomic landscape and known driver gene landscape for OAC^11, 31^ **(Figure 5, A and B, Supplementary Figures 9 and 10)**. Mirroring prior studies from patient samples^11, 31^, alterations in PDOs were enriched for copy-number alterations (21.4%) rather than point mutations or small insertion-deletion mutations, even using stringent thresholds for amplifications and deletions. As with patient series, *TP53* was the most frequently altered gene (73%) in the profiled PDOs, followed by *ARID1A* (24%), *MUCc* (22%), *CDKN2A* (20%), *KRAS* (20%), and *PIK3CA* (17%), although again concordance with matching tissues varied and no PDO *TP53* mutation was identified in 16% (n=4/25) of cases in which a mutation was present in matched tissues **(Figure 5, A and C, Supplementary Figure 10)**.

**Figure 5:**
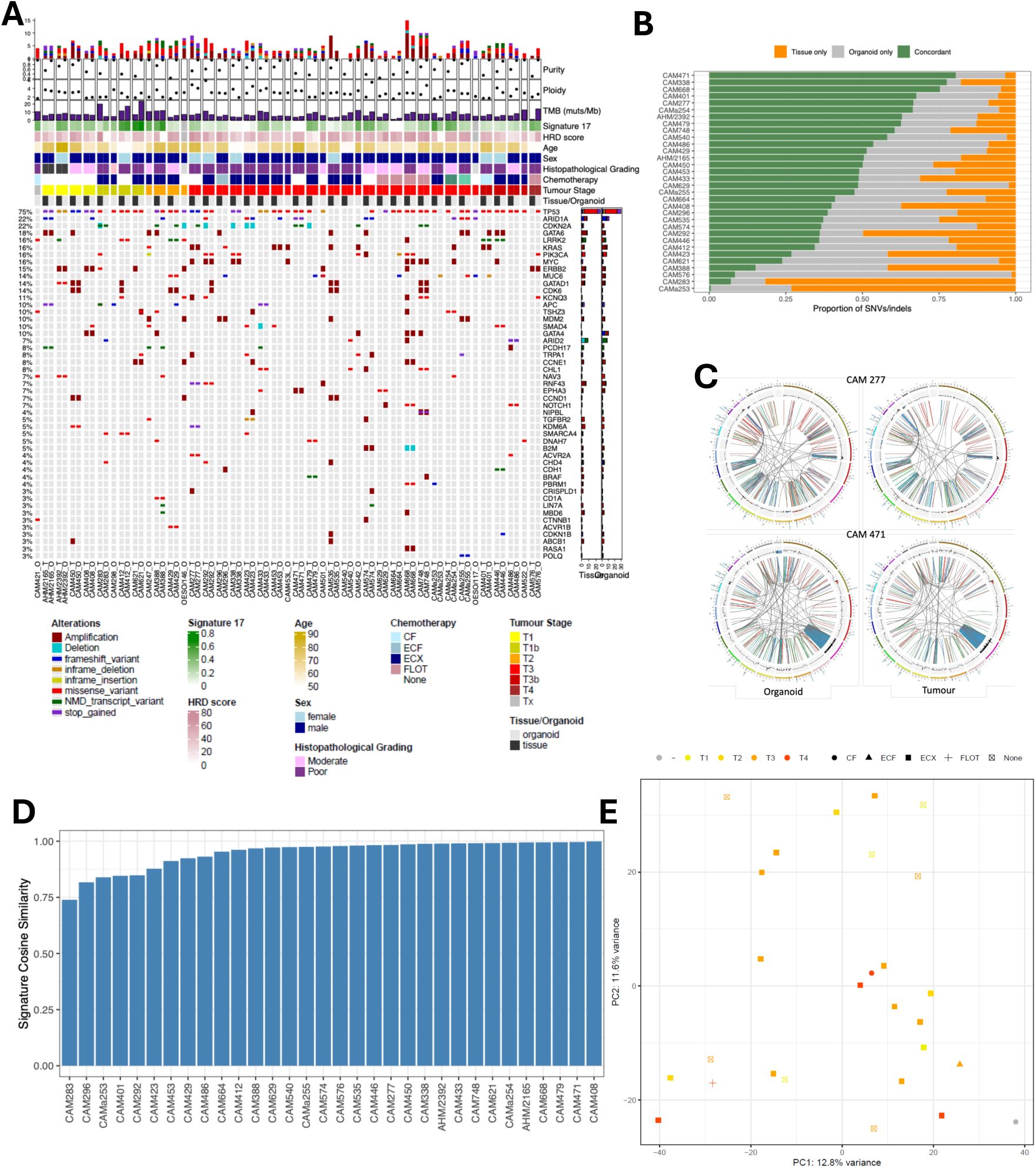
An OAC PDO biobank captures parent tissue features and recapitulates multi-level population diversity. **(A)** Driver alterations and genomic features across OAC-derived PDOs and adjacent tissues. Driver genes were considered according to those previously defined by Frankell et al^11^. **(B)** Concordance of mutational profiles across OAC tissue-PDO pairs, defined by the maproportion of mutations that are shared or unique across samples. **(C)** Circos plots demonstrating genomic features of OAC PDOs and their parent tissues. **(D)** Comparisons of mutational signature profiles across tissue-PDO pairs calculated by cosine similarity. **(E)** Principal component analysis of bulk RNAseq profiles of derived OAC PDOs and relationship to tumour stage.

As in BO, there was broad alignment in the mutational signature landscape of PDOs and matching tissues(median cosine similarity 0.98, range 0.74-1.00; **Figure 5D**). All previously reported^24^ OAC signatures (SBS17a/b, SBS18, SBS40, SBS1 and SBS5) were present **(Supplementary Figure 9I),** with strong PDO–tissue concordance in the BO and OAC-defining^24, 25^ SBS17a (r=0.93, p<0.001) and SBS17b (r=0.92, p<0.001) signatures **(Supplementary Figure 9J)**. Compared with their premalignant equivalents, OAC PDOs displayed increased SBS17a/b, SBS40, and marginal increases in APOBEC-related SBS2/13 **(Supplementary Figure 9K)**. The OAC PDO transcriptomic profile was variable, though there we did not observe specific gene expression signatures that distinguished different groups of PDOs **(Figure 5E, Supplementary Figure 9, L and M).** We also observed recurrent, ploidy-adjusted copy number gains (≥2 relative to sample ploidy) in key oncogenes, including of the receptor tyrosine kinase HER2 in three PDOs, with related protein overexpression **(Supplementary Figure 11).**

To compare OAC PDOs with BO growth metrics, we again used high-resolution time-lapse microscopy **(Supplementary Figure 12, A to C)** Unlike BO, organoids with a solid structure represented 99% of OAC PDOs observed at all assessed timepoints **(Supplementary Table 9)**. Single cells contributing to OAC PDOs were significantly larger than those forming BO PDOs **(Supplementary Figure 12D)** but demonstrated a similar growth rate **(Supplementary Figure 12E)**. A majority of PDOs grew in the seven days after seeding and did so with minimal underlying variation in their growth rates **(Supplementary Figure 12, F and G)**, while some organoids more rapidly increased in size because of fusion events **(Supplementary Figure 12H)**.

### OAC PDOs reproducibly show variable responses to anti-cancer therapies in keeping with their molecular and phenotypic heterogeneity

For OAC a key question is whether PDO phenotypic and molecular diversity translates to differences between PDOs in their sensitivity to specific forms of systemic anti-cancer therapy as well as EBRT, and whether the individual characteristics of each PDO result in differences in sensitivity to individual chemotherapeutics, targeted agents and EBRT. To assess this, five PDOs derived from treatment-naïve donors (OESO117, OESO146, CAM277, CAM408, CAM574) were first exposed to the microtubule inhibitor (docetaxel), anti-metabolite (5-fluorouracil (5-FU), with leucovorin) and platinum (oxaliplatin) constituents of the current perioperative standard-of-care cytotoxic regimen, FLOT **(Figure 6A, Supplementary Table 10)**. Mean IC_50_ values were lowest for docetaxel and demonstrated minimal variation between studied PDOs (1.64–3.04 nM), whereas IC₅₀ values for 5-FU/leucovorin (8.31–51.33 µM) and oxaliplatin (IC₅₀ 7.73–96.78 µM) were higher and demonstrate substantial variation. In contrast, 5FU/leucovorin E_max_ values were 93% or above in all but two PDOs but varied between 62.89-95.82% for oxaliplatin, and between 50.04-93.53% for docetaxel. Validating the utility of using PDOs as a pharmacokinetic model, there was no correlation between IC₅₀ and E_max_ values for 5-FU/leucovorin (r=0.10, p=0.87), oxaliplatin (r=-0.83, p=0.085) or docetaxel (r=0.51, p=0.38). We also demonstrated a heterogeneity of response to external beam x-ray irradiation **(Figure 6B, Supplementary Table 12)**. Organoid forming efficiency varied between 0.42-1.03 in response to a standard 2Gy single fraction of EBRT, and between 0.24-0.75 in response to a high 8Gy single fraction. This corresponded to an integral area under the curve (AUC) for response to 0-8Gy single-fraction irradiation of between 0.46-0.98.

**Figure 6:**
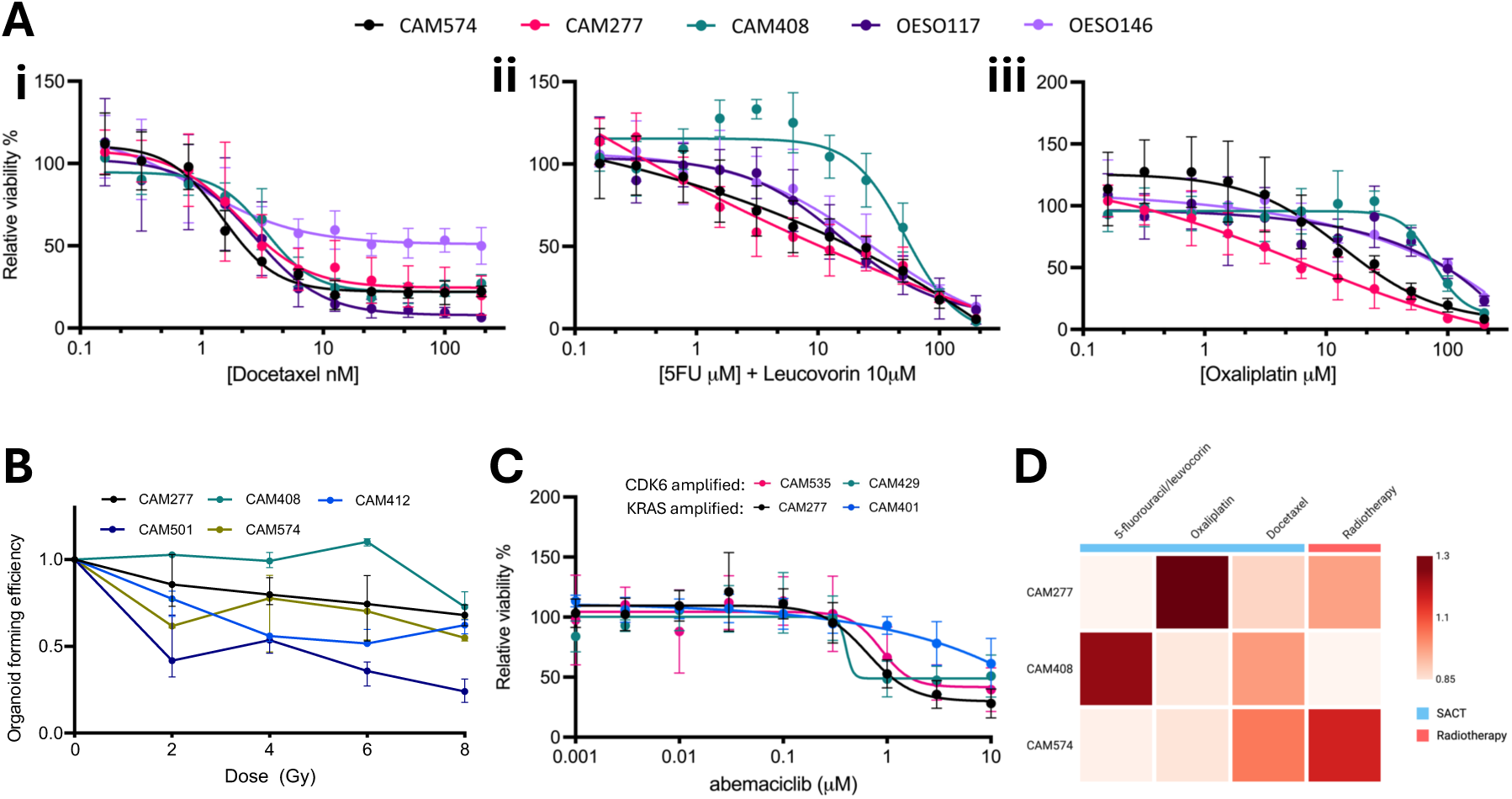
OAC PDOs demonstrate variable responses to chemotherapy, radiotherapy and targeted agents. **(A)** Inhibitory concentration curves demonstrating the viability of five OAC PDOs following exposure to **(i)** docetaxel, **(ii)** 5-fluorouracil (5-FU) and leucovorin, or **(iii)** oxaliplatin. **(B)** Organoid forming efficiency following exposure to a single fraction of 0-8Gy x-ray irradiation for eight OAC PDOs. **(C)** Correlation between AUC organoid forming efficiency following x-ray irradiation and AUC viability values following exposure to 5-FU and leucovorin, oxaliplatin or docetaxel. **(D)** Inhibitory concentration curves demonstrating he viability of four OAC PDOs following exposure to the CDK4/6 inhibitor, abemaciclib. **(E)** Normalised AUC sensitivity values for each anti-cancer treatment across threeseparated PDOs (created in https://BioRender.com).

We and others have also previously reported that as many as 50% of cases of OAC contain sensitising events for CDK inhibitors^11^. However, the extent to which this translates to a consistent therapeutic response across an otherwise heterogenous population is uncertain. Using four PDOs with relevant vulnerabilities (CAM277, CAM401, CAM429, CAM535), we demonstrate diverse sensitivity to the CDK4/6 inhibitor abemaciclib (IC₅₀ 0.57–4.17 µM, Emax 38.8–71.99%) **(Figure 6C, Supplementary Table 13)**. Activity was greatest in CAM277 (KRAS ecDNA, IC₅₀ 0.78 µM, Emax 71.99%) and CAM535 (CDK6-amplified 0.80 µM, 60.37%), intermediate in the CDK6-amplified PDO CAM429 (0.57 µM, 49.14%), and weakest in CAM401 (KRAS nuclear amplified; 4.17 µM, 38.8%) exemplifying the importance of being able to match the PDO specific genomic profile with the therapy response. As outlined in **Figure 6D**, individual PDO sensitivity to different forms of cytotoxic chemotherapy and to EBRT varied. Overall, chemotherapy sensitivity did not predict radiotherapy sensitivity.

## DISCUSSION

Inter- and intra-lesional heterogeneity are fundamental, hard-to-study barriers to the effective prevention and treatment of OAC^13^. Here, we derive and comprehensively characterise a biobank of 97 bulk PDOs and 28 sc-organoids that recapitulate disease heterogeneity in BO and OAC, including within lesions as well as across patient populations. As summarised in **Table 1**, we demonstrate that the specific position each PDO has in the malignant spectrum cannot be discerned from a single feature and nor can it be assumed from the characteristics of the matching tissue due to the extreme heterogeneity. Instead, the functionally relevant disease state of a given PDO can be deduced from a multi-level assessment of genomic, transcriptomic, protein and morphological characteristics **(Supplementary Figure 13)**. In this manuscript we also provide tools for supporting the quantitative characterisation of PDOs at the molecular and phenotypic level, and demonstrate that sc-organoids help capture subclonal heterogeneity and allow for the maintenance of high-risk populations that are otherwise lost over serial passages. Further, we show that individuals PDOs demonstrate variable sensitivity to EBRT and different forms of systemic anti-cancer therapy, and that responses to these treatments vary across the PDO population. In so doing, we demonstrate the utility of PDOs for studying high-risk disease features in the context of premalignant disease, as well as for analysing variable treatment responses in cancer. Previous attempts to model BO neoplastic progression have relied on animal models and xenografts. Genetic mouse models have provided valuable insights and recapitulate certain features of BO and OAC^32^. However, mice do not develop true intestinal metaplasia with goblet cells. Cancer development in these models also typically requires experimental gene knockout, bypassing the complex genomic and epigenomic landscape that characterises carcinogenesis. It is also challenging and resource-intensive to capture the high degree of inter-patient heterogeneity that is a hallmark of BO and OAC using animal models. We therefore view PDOs, which are in theory more scalable, as complementary to animal models. PDOs have been used to study carcinogenesis in other contexts, such as for colorectal cancer^33,34^. However, establishing and propagating precancerous PDO cultures remains a long-recognised challenge, with reported success rates between 0-70%^32^, and the studies in this area remain limited (**Supplementary Table 14**).

**Table 1:** Integrated framework for classifying Barrett oesophagus and oesophageal adenocarcinoma organoids. A decision-based framework to distinguish normal gastric and precancerous organoids are also described in Supplementary Figure 14.

| **Features / dimensions** | **Normal gastric organoids** | **Non-dysplastic BO organoids** | **Dysplastic BO organoids** | **OAC organoids** |
|---|---|---|---|---|
| **Driver gene mutations** | No enrichment for OAC-selected driver events; absence of TP53 and CDKN2A alterations typical of Barrett progression | Frequent *CDKN2A* or ARID1A alterations; TP53 generally absent or subclonal | *TP53* alterations common but could be absent; *CDKN2A* or *ARID1A* frequently co-altered, or progression-associated events | High frequency of *TP53* mutations; recurrent copy number alterations affecting *CDKN2A, ARID1A, GATA6, SMAD4* and others |
| **Genome-wide features** | Very low mutation burden; no evidence of chromosomal instability or whole-genome doubling | Intermediate mutation burden (usually > 2/Mb); mutational signatures concordant with Barrett tissue even when SNV overlap is low | Higher mutation burden (usually > 4/Mb); increased genomic instability | Copy-number alterations dominate; high chromosomal instability and frequent whole-genome doubling |
| **Transcriptome and lineage state** | Strong gastric lineage program (e.g. gastric mucins, digestive enzyme genes); minimal intestinal signature. Separated from BO organoids in PCA analysis | Mixed gastric–intestinal state; inverse relationship between gastric and intestinal programs; Separated from gastric organoids in PCA analysis | Separated from gastric organoids in PCA analysis; increased expression of *HOX* genes relative to non-dysplastic BO | Diverse tumour transcriptional states; no single uniform OAC program |
| **Morphology and growth kinetics** | Benign glandular morphology; used as baseline rather than growth comparison | Four morphologies: single-layer single-lumen; single-layer multi-lumen; multi-layer luminal; solid. Luminal types expand more robustly | Loss of single-layer multi-lumen morphology; increased proportion of solid organoids | Predominantly solid morphology; cell fusion events contribute to growth variability |

Here, we demonstrate derivation of 21 PDOs from regions of non-dysplastic and dysplastic BO with success rates of 68% and 61%, respectively. All remained viable over multiple passages, and all were successfully recovered from cryopreservation. This contrasts with our previous unsuccessful attempts to derive BO PDOs^16^, which likely reflects the addition to culture medium of gastrin and FGF10^32^. There is also uncertainty as to the degree to which PDOs can be used to study BO dysplastic progression. Previous attempts to develop a model for OAC carcinogenesis have primarily focussed on gene editing of BO or normal gastroesophageal junctional PDOs^35, 36^. However, genetic manipulation in this context is inefficient and poor survival rates result in selection and propagation of monoclonal populations^35, 36^. Our work and that of others has also shown that OAC very likely always derives from BO^5, 37^, which questions the relevance of modelling dysplastic progression using normal gastroesophageal junctional PDOs. To this end, studies in colorectal cancer have demonstrated that modelling tumourigenesis using normal tissue PDOs as a starting point fails to capture many of the known molecular changes that accompany the development of invasive disease^33, 34^. There is therefore a need to model progression using PDOs derived from premalignant tissue.

Our comprehensive genomic, transcriptomic and phenotypic characterisation of BO PDO cultures demonstrates that whilst PDOs derived from dysplastic tissue exhibited more cancer-like genomic alterations than those derived from non-dysplastic tissue, they have lower concordance with matching tissue. Concordance with matching tissue was also generally lower for BO than it was for OAC PDOs. This is likely to be due to the nature of heterogeneity in BO, which is characterised by the presence of distinct clones that do not sweep the entire segment and instead arise through clonal expansion that occurs on a scale of millimetres^7, 26^. Although morphological analyses were performed on a limited number of representative PDO lines, the observed structural differences are consistent with established hallmarks of polarity loss in dysplastic progression. However, further experimental work is required to provide direct evidence for the underlying mechanisms, and organoid models represent an excellent platform for such future investigations.

We reasoned that it is also possible that high risk subclonal populations may not be adequately reflected in bulk PDO sequencing and may be lost over time or through passaging. To counter this, we demonstrate a technique to isolate sc-organoids, which we show can isolate high risk subclones that may not be represented in bulk PDO sequencing. These can then serve as observable functional proxies for intralesional heterogeneity at a single clone level, with each demonstrating varying degrees of malignant potential. Looking ahead, integrating these clones into epithelial-fibroblast assembloid systems, and incorporating immune cells, such as autologous or allogeneic tissue-resident myeloid or T-cell populations, and immunogenic cytokines into this platform would allow interrogation of how epithelial and stromal evolution interfaces with emerging immune microenvironments during early cancer development and progression^17^.

scRNA-seq of PDOs further reinforced their capacity to recapitulate disease stage-specific cellular heterogeneity, with the abundance of specific cell clusters shifting in a disease stage - dependent manner. Metaplastic Progenitor cells were enriched in non-dysplastic BOPDOs, while Metaplastic Intestinal and Metaplastic Cycling populations were significantly enriched in dysplastic PDOs, the latter associated with higher mutational burden and greater driver mutation load. Together with the broader transcriptomic and genomic data presented here, these findings suggest that increased proliferative and intestinal metaplastic commitment is a hallmark of progression towards dysplasia and early cancer, and provide a functional platform for investigation of how mutational accumulation drives metaplastic lineage evolution.

As well as being able to recapitulate the substantial heterogeneity in OAC organoids, we also show that we can recapitulate the diverse treatment responses seen in patients with OAC, demonstrating highly variable responses to each of the individual components of the standard preoperative regimen, FLOT^14^. There is, for example, consistently high sensitivity (i.e. low IC_50_ values) to docetaxel, though this has a variable maximum effect (i.e. E_max_). In contrast, PDOs have a lower and more variable sensitivity to 5-FU/leucovorin, though this consistently exerts a high maximal effect when added to PDOs at a sufficiently high concentration. This mirrors clinical data, where 5FU/leucovorin is the backbone of most chemotherapy regimens in OAC due to a conserved effect across the entire patient population^38^. It is similarly interesting that chemotherapy sensitivity does not mirror radiation or CDK inhibitor sensitivity, which provides biological rationale for an expanding portfolio of studies that are combining intensified preoperative chemotherapy with chemoradiotherapy or targeted agents. It would be of interest to determine whether the application of the PDOs described here to an organ-on-chip approach, such as that described by *Pal et al*^20^, would further improve modelling of therapeutic response.

In summary, we have developed a comprehensively characterised biobank of BO and OAC PDOs that reflect known disease heterogeneity across pre-malignant and invasive disease. Careful characterisation is essential to understand the provenance of a given organoid and derivation of sc-organoids allows for the analysis of high-risk clones. Together, these models provide a scalable platform for functional studies of OAC carcinogenesis and therapeutic response.

## ACKNOWLEDGEMENTS

This work was supported by grants from Cancer Research UK (CRUK) to L.Z.(A31116 and EDDPJT-May23/100046) and C.M.J. (RNPSF-Jan23/100006), from the Academy of Medical Sciences to C.M.J. (SGL030\1032), and by a Medical Research Council Programme Grant awarded to R.C.F. (MR/W014122/1). The Oesophageal Cancer Clinical C Molecular Stratification (OCCAMS) study and its successor OCCAMS 2 study were funded by Programme Grant funding (A15874, A22720, A22131) from CRUK. D.P.M. is supported by a Clinical Research Training Fellowship supported by the CRUK Cambridge Centre (SEBCATP-2024/100008). C.M.J. was supported for the duration of this work by a Clinical Lectureship partly funded by CRUK RadNet Cambridge (C17918/A28870), a CRUK Clinician Scientist Fellowship (RCCCSF-May26/100001) and a Loke Medical Sciences Fellowship funded by King’s College, Cambridge. The authors are grateful to the Human Research Tissue Bank at Addenbrooke’s Hospital, Cambridge University Hospitals NHS Foundation Trust, and for the support provided to it by the National Institute for Health Research (NIHR) Cambridge Biomedical Research Centre (BRC-1215-200014). The views expressed are those of the authors and not necessarily those of the NIHR or the Department of Health and Social Care. The authors are grateful for the materials provided by the Jacqueline Shields Group, King’s College London, and the Fiona Gribble Group, University of Cambridge.

## AUTHOR CONTRIBUTIONS

R.C.F., C.M.J. and L.Z. conceived the manuscript, obtained funding and oversaw the work. C.M.J. conceived therapeutic analyses in OAC PDOs. L.Z. conceived studies relating to Barrett oesophagus PDOs and single cell-derived clonal organoids. D.H.J., G.D. and SG.J. conducted bioinformatic and computational analyses. D.P.M., Y.W., C.C., T.S.C., H.C., X.L., A.M., C.M.J. and L.Z. performed wet laboratory experiments and led or contributed to experimental design. Y.W. generated the scRNA-seq dataset and performed initial analysis. E.B. and B.H. conducted the in-silico analysis of organoid morphologies. K.T.M., M.dP. and K.S-P. contributed to sample collection, curation and characterisation. D.H.J., D.P.M., R.C.F., C.M.J. and L.Z. wrote the first draft of the manuscript. All authors contributed to subsequent revisions and all have read and approved the manuscript.

## METHODS

### Study approach and regulatory approval

Samples were obtained from surgical or endoscopic resection specimens obtained from patients with known BO, dysplasia or OAC (sampling and PDO biobank summarised in **Supplementary Figure 1)**. Prior ethical approval was provided by the United Kingdom Health Research Authority, including: “A study to investigate the cell biological determinants of the development and progression of the pre-malignant condition in Barrett’s oesophagus” (Biomarker), 01/149; “Multicentre Study to determine predictive and prognostic biomarkers and therapeutic targets for oesophageal and junctional adenocarcinoma including whole genome sequencing” (OCCAMS), 10/H0305/1; “A study using tissue to develop stem cell-based therapies and to investigate human physiology, pathology and embryonic development” (Organ donor), 15/EE/0152. All patients provided written informed consent prior to sampling, which was performed within a National Health Service setting.

### Sampling strategy

To derive non-dysplastic BO PDOs two to three biopsy specimens were taken from the same region at surveillance endoscopy, with a preference for adjacent sampling wherever possible. All dysplastic BO PDOs and a single OAC PDO were obtained from specimens taken via a punch biopsy from an endoscopic mucosal resection (EMR). For OAC models, resection specimens were evaluated and samples obtained for organoid derivation and adjacent tissue for snap freezing. For control gastric PDOs and intestinal PDOs, both patients and disease-free organ donor tissues were used. Samples were briefly minced and mixed to ensure homogeneity, and half or two thirds of the tissue mixture was immediately processed to derive PDOs, the methods for which are outlined below. The remnant was snap-frozen using liquid nitrogen and stored at - 80°C for profiling of adjacent tissues recognising that given the heterogeneity there will be potential differences at the cell and molecular level. To further mitigate and understand the genotype to phenotype correlation in all cases, a section was cut from each snap frozen sample and stained with haematoxylin and eosin (HCE) prior to DNA extraction. The HCE-stained slide was independently reviewed by at least two experienced pathologists to assess cellularity, and to confirm malignancy for OAC or to assess for the presence and severity of dysplasia in BO. In all instances, only samples with at least 70% cellularity were used for deoxyribonucleic acid (DNA) or ribonucleic acid (RNA) extraction using the AllPrep kit (Qiagen). DNA was extracted from blood samples using the QIAmp DNA Blood Maxi kit (Qiagen). In all cases a germline reference was obtained from blood wherever possible or otherwise from distant normal mucosa.

### Derivation of BO PDOs

To derive BO PDOs, biopsy samples were rinsed in ice-cold phosphate buffered saline (PBS) and pooled. These were then minced and digested using recombinant TrypLE Express Enzyme (ThermoFisher) supplemented with 10µM Y-27632 (Tocris Bioscience) at 37°C for up to 10 minutes. Vigorous shaking was used every three minutes to dissociate tissue aggregates. Cells were pelleted at 300g for 5 minutes and resuspended in 150µl TrpLE Express Enzyme. Further mechanical dissociation was achieved using a P200 pipette prior to centrifugation at 300g for 5 minutes. The resultant single cell mixture was resuspended in Cultrex Basement Membrane Extract (BME) Type 2 (RCD Systems) and plated as 15µl domes for ongoing culture.

### Derivation of OAC PDOs

Samples obtained from surgical resection specimens were processed as per our previously reported protocol^16^. Briefly, each sample was rinsed in ice-cold PBS, minced then digested through incubation at 37°C with collagenase II (1.5mg/ml) for up to 2 hours. Large fragments were then removed by filtration using a 70µM cell strainer and the resultant cell suspension centrifuged at 400g for 2 minutes. The pellet was resuspended in PBS as a further wash step to remove debris and a further identical centrifugation step performed. Isolated cells were then resuspended in Cultrex BME Type 2 and plated as 15µl-20µl domes for ongoing culture.

### Derivation of gastric and intestinal PDOs

Gastric PDOs were established either from endoscopic biopsies of the gastric cardia from BO patients or from gastroesophageal junction (GOJ) tissue from disease-free organ donors. For gastric cardia PDOs derived from BO patients, biopsy specimens were processed using the same protocol as for BO PDO derivation. Gastric PDOs from organ donors were generated as described in our previous study^5^. Briefly, a circumferential tissue fragment (up to 10 cm in length) spanning 3–5 cm of distal oesophagus and 3–5 cm of proximal stomach was obtained. GOJ PDOs were established from a 5 mm tissue strip spanning the Z-line (squamous–columnar junction). Gastric body PDOs were derived from tissue located at least 2 cm below the Z-line. Intestinal PDOs were kindly provided by Professor Fiona Gribble, University of Cambridge. In brief, intestinal organoids were generated from healthy jejunal or duodenal tissue obtained from patients undergoing gastrectomy or pancreaticoduodenectomy **(Supplementary Table 2)**. Note: although organ-donor-derived organoids are not strictly patient-derived, we refer to them as PDOs for consistency and ease of interpretation throughout the manuscript.

### PDO culture

All PDO cultures were maintained in a humidified atmosphere with 5% carbon dioxide (CO_2_) at 37°C. PDOs were maintained in IntestiCult™ (INC) Organoid Growth Medium (StemCell Technologies) supplemented with Primocin (1mg/ml, InvivoGen) and Y-27632 (10µM), with or without FGF10 (100ng/ml, Peprotech) and Gastrin (10nM, Sigma). Media was refreshed every two to five days for cell maintenance outside of specific experimental conditions. PDOs were passaged at least every two weeks through the addition of pre-warmed TypLE Express Enzyme and manual dissociation by pipetting. The resultant cell solution was centrifuged at 300g for 5 minutes, providing a cell pellet that was resuspended in Cultrex Basement Membrane Extract, Type 2 and plated as 15µl-20µl domes that were allowed to polymerise prior to the addition of media. Once established, BO and intestinal PDO cultures were maintained in organoid medium (OWR) prepared from Advanced Dulbecco’s Modified Eagle Medium (DMEM)/F-12 supplemented with 0.5nM Wnt Surrogate-Fc Fusion Protein (N001, U-Protein Express BV), 9.6nM RSPO3-Fc Fusion Protein conditioned medium (R001, U-Protein Express), 10mM HEPES, 1× Glutamax, 1× N2, 100ng/mL Noggin, 10 mM Nicotinamide, 1 mM N-acetylcysteine, 1×B27, 50ng/mL EGF, 500nM A8301, 10μM SB202190, 1× Primocin, 10nM Gastrin, 10μM Y-27632 and 100ng/mL FGF10.

### Establishment of single cell-derived clonal organoids

Sc-organoids were microscopically isolated and collected from P0 bulk PDO cultures, typically five to six days after seeding, when the cultures had reached approximately 70–80% confluency. Briefly, culture medium was removed and replaced with an ice-cold mixture comprising of a 1:1 ratio of TrypLE Express Enzyme and Advanced Dulbecco’s Modified Eagle Medium (DMEM)/F-12 supplemented with 10µM Y-27632 (TrypLE/ADF). A P1000 pipette was then used to manually break the BME Type 2 domes whilst leaving PDOs intact. The resultant mixture was transferred to the underside of a sterile Corning cell culture plate lid with multiple 40µl TrypLE/ADF droplets. Using a brightfield microscope, the PDO suspension was inspected for single organoids exceeding a diameter of 200µm. Once identified, these were aspirated under direct vision using a P20 pipette and transferred to a TrypLE/ADF droplet. This process was repeated multiple times and each droplet then transferred to a fresh 1.5ml microcentrifuge tube. This was incubated at 37°C for five minutes using a ThermoMixer® (Eppendorf). A P20 was then used to mechanically dissociate the isolated PDO and the resultant single cell solution resuspended in Cultrex BME Type 2 and plated as 15µl domes for ongoing culture. Of note, to study heterogeneity in BO and dysplasia, single organoids were picked from passage 0 cultures, i.e. cultures established directly from tissue without any passaging, to ensure that distinct clonal populations were selected.

### PDO treatment using systemic anti-cancer therapies

The viability of OAC PDOs was assessed following exposure to individual components of FLOT (all supplied by SelleckChem) and to the cell cycle inhibitor, abemaciclib (LKT labs). To mirror clinical administration, oxaliplatin and docetaxel were administered to PDOs as single agents whereas 5-fluorouracil was administered in combination with a fixed dose of leucovorin. Each drug was reconstituted as per manufacturer instructions and each was diluted in either DMSO (5 - fluorouracil, docetaxel, and abemaciclib) or ultrapure water (oxaliplatin and leucovorin) to values within a range of 200µM-0.195µM for components of FLOT, except for leucovorin, which was administered at 10µM on all occasions, and 0.001-10μM for abemaciclib. Values for both dilution series were selected based on others reported in the literature^39, 40^. The concentration of DMSO was fixed at ≤0.1% within treatment and control vehicle wells.

In line with existing protocols, PDOs were dissociated using TrypLE Express Enzyme then counted and resuspended in BME Type 2 prior to drug exposure^16, 41^. A total of 5000 cells were seeded within 20µL BME Type 2 droplets formed at the centre of each well of a 96-well plate. These were allowed to polymerise prior to the addition of media. Cultures were maintained for 72 hours and then exposed to drug when in a logarithmic growth stage. Viability was assessed after 144 hours exposure to components of FLOT, abemaciclib, or vehicle. Media, including drug or vehicle, was refreshed after 72 hours for PDOs treated with FLOT constituent drugs.

### PDO treatment using external beam radiotherapy

Prior to irradiation, established PDOs were dissociated to a single cell solution. This was performed by first dissociating PDOs from Cultrex Basement Membrane Extract, Type 2, through incubation with pre-warmed TypLE Express Enzyme and manual agitation by pipetting. Dissociated PDOs were pelleted by centrifugation at 300g for 5 minutes and resuspended in 30µl cool PBS. A P20 pipette was then used to manually dissociate the PDOs to a single cell solution and the cell concentration assessed using a Countess II Automated Cell Counter (Invitrogen). A total of 10,000 cells were plated in 15µl Cultrex Basement Membrane Extract, Type 2 with media added once this had polymerised. The media was replaced with pre-warmed PBS prior to irradiation 24 hours later using a CIX1 (XStrahl) benchtop irradiator. Single fraction treatments of 0Gy (sham), 2Gy, 4Gy, 6Gy and 8Gy x-ray irradiation were delivered and PBS then replaced with media 15 minutes after irradiation.

### Whole genome sequencing

Extracted DNA was subjected to paired-end whole genome sequencing at a depth of 50x for tissues, 30x for PDOs and 15x for sc-organoids, using a well-established pipeline^11^. A combination of in-house tools and FastQC (http://www.bioinformatics.babraham.ac.uk/projects/fastqc) were used for quality control. Sequencing reads were aligned against the hg19/Ensembl GRCh37 reference genome for mutation calling using the Burrows-Wheeler BWA-MEM alignment algorithm^42^. Aligned reads were subsequently sorted into genome coordinate order and Picard (http://broadinstitute.github.io/picard) used to remove duplicate reads. Single nucleotide variants and insertion deletion mutations were called with Strelka v2.0.15^43^. Tumour purity and ploidy were inferred with ASCAT-NGS v2.17, and copy number alterations derived with ASCAT from read counts at germline heterozygous sites identified by GATK HaplotypeCaller v3.2-2, adjusting for the estimated normal-cell component. Structural variants (SVs) were called using MANTA v0.27 with any SVs that had any supporting reads in the matched normal being discarded as previously described^44^.

### Targeted amplicon sequencing

Targeted sequencing was performed using the IDT xGen™ Amplicon Core Kit according to the manufacturer’s protocol, with a custom panel targeting driver mutations identified by WGS in Figure 2, including *CDKN2A, MUCc, ZFHX3, PTEN, PCDH17, ARID2, SMARCA4,* and *TP53*. For matching tissue and the bulk PDO, 3.6 and 2.3 ng of DNA were used, respectively, due to the limited amount of remaining sample material. Sequencing reads were aligned against the hg38 reference genome using the BWA-MEM alignment algorithm as specified earlier, and variants were called using Strelka v2.9.10. Specific variants were mapped across from hg19 to hg38 using https://liftover.broadinstitute.org/.

### RNA sequencing

Extracted RNA was quality assessed using the RNA 6000 Nano Kit (Agilent) on a 2100 Bioanalyser (Agilent). Samples for which there was sufficient material available for further analysis and that had a calculable RNA integrity number were determined to have passed quality control and were quantified using the Qubit High sensitivity RNA assay kit (Thermo Fisher). The TruSeq Stranded Total RNA High Sensitivity protocol with ribosomal depletion was used to prepare libraries using an input of at least 100ng but preferably 250ng if available. KAPA quantification (KAPA Biosystems) was used to quantify libraries, which were pooled and run on a HiSeq 4000 to generate 75 base pair adjacent-end reads. After sequencing, reads were aligned by STAR. Gene-level counts for each sample were generated using Bioconductor’s Genomic Alignments package together with ENSEMBL gene annotations. These raw counts, along with sample-specific sequencing depths and gene lengths, were then used to calculate Transcripts Per Kilobase Million (TPM) values for individual genes across all samples.

### Single cell RNA sequencing

Single cell suspensions were prepared from organoids after 7 days culture for submission to the 10x Genomics 3′ Gene Expression v4 4-plex On-Chip Multiplexing workflow. Cells were washed with cold PBS to remove residual extracellular matrix and transferred to 15 mL centrifuge tubes and dissociated in pre-warmed TrypLE with Y-27632 at 37°C with gentle trituration until predominantly single cell suspensions were obtained. Dissociated cells were centrifuged at 300 × g for 5 min at 4°C, resuspended in cold PBS containing 0.4% BSA, and passed through 40 μm cell strainers. Filtered suspensions were centrifuged again at 300 × g for 5 min at 4°C. Dead cells were removed using the Dead Cell Removal Kit (130-090-101; Miltenyi Biotec) and MS columns according to instructions. The live-cell fraction was washed with PBS containing 0.04% BSA, centrifuged at 300 × g for 5 min, and resuspended in PBS containing 0.04% BSA. Cell concentration and viability were determined using a NucleoCounter NC-202. Samples with minimal debris, low aggregate levels, and viability above 70% were taken forward. Samples were maintained on ice before proceeding with the 10x Genomics protocol. Reads were processed using CellRanger with alterations for on-chip multiplexing, as integrated through Nextflow^45^.

### Bioinformatic analyses

In determining relevant driver events, we considered mutations in 76 genes identified by Frankell et al. as positively selected for in OAC^11^. Amplification events were defined as cases where copy number was more than double the sample ploidy, whilst deletion events were strictly defined as a total copy number of zero. Losses in the single cell-derived clonal organoids were defined as a minor copy number of zero. Two PDOs (OESO/117 and OESO/146) were used for chemotherapy treatment screens but underwent WGS elsewhere. Clinical and genomic information for these PDOs was obtained from Cell Model Passports (https://cellmodelpassports.sanger.ac.uk/).

The HRD index score, the sum of three HRD-associated copy number features applied here as a broad measure of total chromosomal instability, was calculated using the scarHRD R package^46,47^. Mutational signatures were extracted de novo using SigProfiler applied to the complete set of OAC tissues and PDOs, resulting in the following signatures: SBS1, SBS2, SBS5, SBS8, SBS13, SBS17a, SBS17b, SBS18, SBS31, SBS35, SBS40, SBS93^48^. These signatures were then fitted using the deconstruct Sigs R package with a 2% prevalence cut-off^49^. Signatures SBS31 and SBS35 were not fitted for BO PDOs since they are caused by platinum chemotherapy. Subclonal reconstruction of the non-dysplastic BO PDOs was conducted with PyClone-VI using the beta-binomial distribution^50^.

Phylogenetic analysis of the single cell-derived clonal organoids was conducted using the parsimony ratchet method as implemented through the phangorn R package^51^, with branch lengths assigned according to the ACCTRAN criteria. Specific SNVs were assigned to branches onto which they fitted exactly based on their presence across organoids deriving from the respective branch. A subset of the above signatures (SBS1, SBS2, SBS5, SBS8, SBS13, SBS17a, SBS17b, SBS18, SBS40) with established presence in BO, and that do not relate to chemotherapy treatment, were fitted using deconstruct Sigs to any branch with more than 25 unique SNVs. Barplots for signature contributions were plotted onto the corresponding branch and extended manually to fit the branch length.

Differential gene expression analysis was conducted using the DESeq2 R package^52^. Functional enrichment of differentially expressed gene sets was analysed using the gProfiler2 R package^53^, specifically on genes with p-value (false discovery rate adjusted for multiple testing) of less than 0.05 and with an absolute log2-fold change greater than 1, and separated into up- and down-regulated genes. This package enables enrichment analysis against multiple databases including Gene Ontology, the Human Protein Atlas for cell-type analysis, and TRANSFAC for transcription factor-binding profiles. Gastric and intestinal gene set ‘scores’ were calculated through gene set variation analysis of the TPM-normalised data, using the GSVA R package^54^, taking *MUCc*, *MUC5AC*, *CLDN18*, *PGC*, *GKN1*, *GKN2* as gastric genes and *MUC2*, *TFF3*, *REG4*, *GPA33*, *CDX2*, *FABP2*, *OLFM4*, *ANPEP* as intestinal genes.

RNA-seq counts for OAC-derived PDOs underwent batch correction using ComBat, as part of the sva R package^55^. Principal component analysis was conducted on the top 1000 variable genes following batch correction and variance stabilising transformation. Gene set enrichment analysis, using the cancer hallmark gene sets, was applied on the loadings of each gene for the first two principal components using fgsea. Normal gastric, normal intestinal, and BO-derived PDOs were sequenced in the same batch and so correction was not required.

For scRNA-seq, cells with fewer than 1,500 unique features and greater than 25% mitochondrial fraction were removed. Normalisation, scaling, cell cycle scoring, and further analysis were conducted using Seurat^56^. Batch correction was conducted using Harmony, following which Find Neighbors was applied to the first 15 principal components and Find Clusters with a resolution of 0.48 was used to identify twelve clusters, as identified as optimal via an Elbow plot ^57^.

### Organoid forming efficiency assay

All PDOs were imaged on the day of, and 10-days following, irradiation using an Incucyte® SX5 Live-Cell Analysis System (Sartorius, Goettingen, Germany). The Incucyte® Organoid Analysis Module (Sartorius, Goettingen, Germany) was used to count the number of PDOs at each assessed timepoint using a segmentation radius of 100µm, segmentation sensitivity of 30, segmentation edge split with sensitivity of 100, cleanup with hole fill of 500,000µm ^2^ and an adjusted pixel size of -1. Filters were set to count objects as PDOs only with a minimum area of 1500µm^2^ and maximum eccentricity of 0.25 as a strict cut-off to exclude budding, migrating or necrotic PDOs. Given that there should not be PDOs formed at day zero, the PDO count at this point was considered to reflect noise and was subtracted from the day 10 PDO count. Organoid formation efficiency was calculated for each radiation dose as the number of formed PDOs at day 10 divided by the cell seeding number, 10,000. Relative organoid forming efficiency was determined by dividing the organoid forming efficiency at each delivered dose by the organoid forming efficiency at 0Gy.

### PDO viability

PDO viability was quantified using a luminescent assay 72- or 144-hours after respective administration of abemaciclib or FLOT. In accordance with manufacturer instructions, 20uL of CellTiter-Glo® 3D (Promega) was applied to each well of a 96-well plate, which was then shaken for five minutes. Plates were then incubated for 30 minutes in a humidified atmosphere with 5% CO_2_ and luminescence read using an Infinite 200 Pro microplate reader (Tecan).

### PDO fluorescent labelling

PDOs derived from BO and OAC were labelled with fluorescent nuclear markers via lentiviral transduction. Briefly, PGK-H2B-eGFP or PGK-H2B-mCherry plasmids were co-transfected with pMD2.G and psPAX2 into HEK293 cells (a gift from Sakari Vanharanta, University of Helsinki) to produce lentivirus, following the protocol described in Kita-Matsuo et al^58^. Viral supernatant was concentrated 16-fold using the Lenti-X™ Concentrator (631231, Takara Bio) and resuspended in organoid medium. Organoids were harvested, resuspended in 15 µL (approximately 500,000 cells), mixed with 250 µL of concentrated lentivirus, and incubated for 4 hours at 37 °C. The organoids were then washed, seeded, and maintained under standard culture conditions.

PGK-H2BeGFP was a gift from Mark Mercola (Addgene plasmid # 21210; http://n2t.net/addgene:21210; RRID: Addgene_21210). PGK-H2BmCherry was a gift from Mark Mercola (Addgene plasmid # 21217; http://n2t.net/addgene:21217; RRID: Addgene_21217). pMD2.G was a gift from Didier Trono (Addgene plasmid # 12259; http://n2t.net/addgene:12259; RRID: Addgene_12259). psPAX2 was a gift from Didier Trono (Addgene plasmid # 12260; http://n2t.net/addgene:12260; RRID: Addgene_12260).

### Fluorescent in situ hybridisation

Flourescent *in situ* hybridisation (FISH) was performed using routine diagnostic *HER2* probes. As previously described^59^, PDO cultures were prepared using standard cytogenic procedures for harvesting, fixation (3:1 methanol : acetic acid solution) and slide formation. All subsequent pre-treatment and hybridisation steps were undertaken in accordance with manufacturer instructions by the Department of Histopathology, Cambridge University Hospitals NHS Foundation Trust.

### Time-lapse imaging of PDO growth and morphology

Time lapse imaging was performed using a Zeiss Axio Observer Live Cell Station at 10x magnification at either 2-4 hourly intervals or at day 4, 6 and 7 after single cell seeding. Up to four locations were imaged at each time interval with images captured across up to 23 slices through a z-stack of the Cultrex BME Type 2 dome in which the PDOs are cultured. Brightfield images were acquired, in addition to GFP fluorescence via excitation at 469nm with emission recorded at 525nm using a GFP filter set, and mCherry fluorescence via excitation at 590nm with emission at 687nm using an mCherry filter set. Exposure time and acquisition parameters were kept constant at all imaging intervals. Images were processed using Zeiss Zen software.

### Computational analyses of PDO growth kinetics and morphology

Once processed, images of PDOs obtained at each timepoint were analysed using the Fiji distribution of ImageJ^60, 61^. Given an inherent partial eccentricity, each PDO was manually delineated using an ellipse. The minimum and maximum widths of this were averaged to give an approximate value for the PDO diameter. Though often visible across multiple levels of a z-stack, PDOs were measured only once, at the plane at which they were most in focus. PDOs were simultaneously categorised by morphology and distinguished by the presence or absence of a visible lumen, which if present was categorised as single or multiple, and whether the cells surrounding the lumen were a single cell layer or of multiple layers. For hollow PDOs (i.e. those with a lumen), the number of cells within that PDO was presumed to be proportional to the square of the diameter (i.e. the surface area). For solid organoids (i.e. those lacking a lumen), the number of cells in that PDO was presumed to be proportional to the cube of the PDO (i.e. the volume of the organoid).

An *in silico* cell-level model of PDO growth was built in CompuCell3D (CC3D) Version 4.2.5^62^. This implements a lattice-based multi-site Cellular Potts Monte Carlo model and has previously been used to model cells growing around a lumen. Briefly, each cell in this model can occupy a number of lattice sites. A model is developed to minimise the energy of the lattice conformation using the following equation:

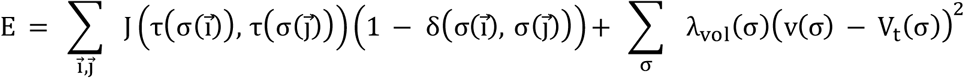

where the first sum represents energy changes due to cell adhesion for every pair of neighbour lattice sites (*i, j)* and the second sum represents volume constraints for every cell, σ. In the first sum, *J* represents different adhesion energies between each pair of cell types 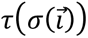 and 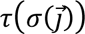, or between a cell type and the extracellular matrix. In the second equation, each cell has a target volume V_t_(σ) and a current volume v(σ). The energy equation penalises deviation from the target volume, with λ_vol_(σ) representing cell elasticity. Multiple Monte Carlo Steps occur, with each considered a time unit proportional to experimental time. At each step each pixel will attempt to copy itself to a neighbouring lattice site. This will succeed with probability 1 if the change reduces the overall energy, E, in the system.

The model was implemented in two dimensions, representing experimentally captured images of PDOs in cross section. Three cell types were used: polar, non-polar and lumen. Each lumen was modelled as a single ‘cell’, with the size of this lumen ‘cell’ determined by the number and type of cells surrounding the lumen.

### Statistical analysis and reproducibility

All experiments were undertaken in at least biological triplicate unless otherwise stated. Given the known challenges of inter-batch heterogeneity in Cultrex BME Type 2 (RCD Systems), all biological replicates for irradiation experiments were performed using the same batch of BME Type 2. Integral area under the curve (AUC) values for organoid forming efficiency following irradiation were calculated using GraphPad Prism Version 10.4.1. for Windows (GraphPad Software, Boston, USA). Percent viability following exposure to systemic anti-cancer therapy was normalised to vehicle control and modelled against drug concentration using a three-parameter logistic (3PL) curve in GraphPad Prism v10.4.1 with Top fixed at 100%, and Bottom and Hill slope free. A four-parameter logistic (4PL) was also fit and retained only if it was clearly supported by the Akaike Information Criterion with small-sample correction (AICc) (ΔAICc ≤ −4) and the top 95% confidence interval excluded 100% across biological replicates; otherwise 3PL was used. Emax is reported as 100 − Bottom and the area under the curve (AUC) was computed over the tested log-dose range and normalised and reported as 1-AUC. Pairwise comparisons were made using the Mann-Whitney U test unless otherwise stated. Comparisons of more than two groups were made using the Kruskal Wallis test unless otherwise stated. Adjustments for multiple significance testing were made using the Bonferroni test unless otherwise stated in instances in which repeated statistical comparisons were drawn from the same dataset.

### Data availability

Metadata to correlate information within this archive to that published here is provided in **Supplementary Table 15**.

### Code availability

The scripts developed during the analysis presented here are available at the following GitHub repository, released under a GNU GPL-v3.0 license: https://github.com/fitzgerald-lab/OAC_Barretts_PDOprofiling. This includes scripts for analysing scRNA-seq data, WGS data (e.g. driver gene profiles, mutational concordance, signature contributions, phylogenetic analysis, clonal dynamic analysis), RNA-seq data (differential expression, unsupervised clustering, pathway and gene-set analysis) obtained from BO, normal tissue-, and OAC-derived PDOs, and sc-organoids.

## LIST OF SUPPLEMENTARY FIGURES

**Supplementary Figure 1:**
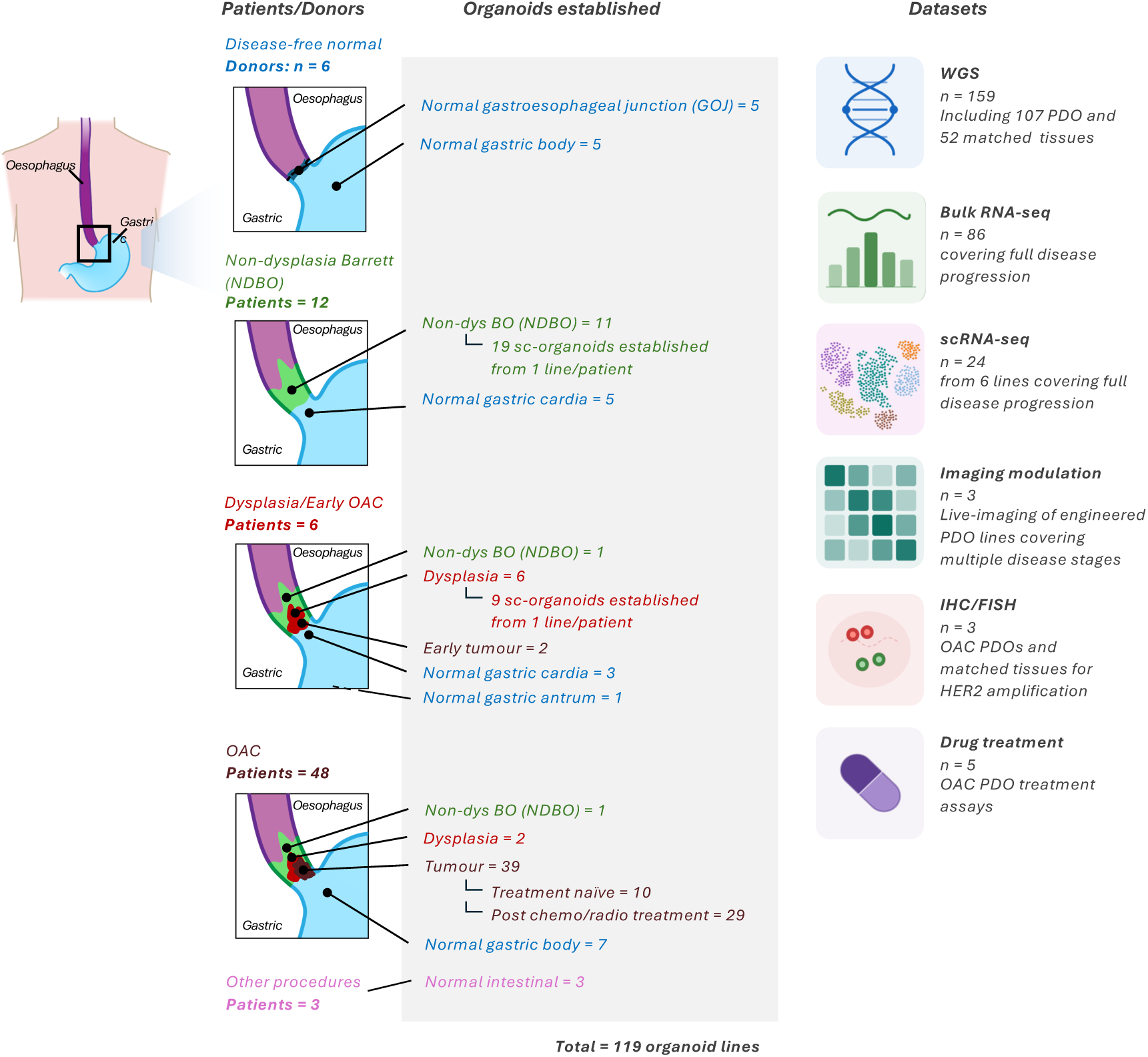
Schematic overview of the 122 normal, Barrett oesophagus and OAC bulk PDOs and single cell-derived organoids generated for this study. The site from which samples were obtained, and the sampling route, is shown, as is a summary of the multi-level characterisation employed for all obtained models. Image generated using BioRender®.

**Supplementary Figure 2:**
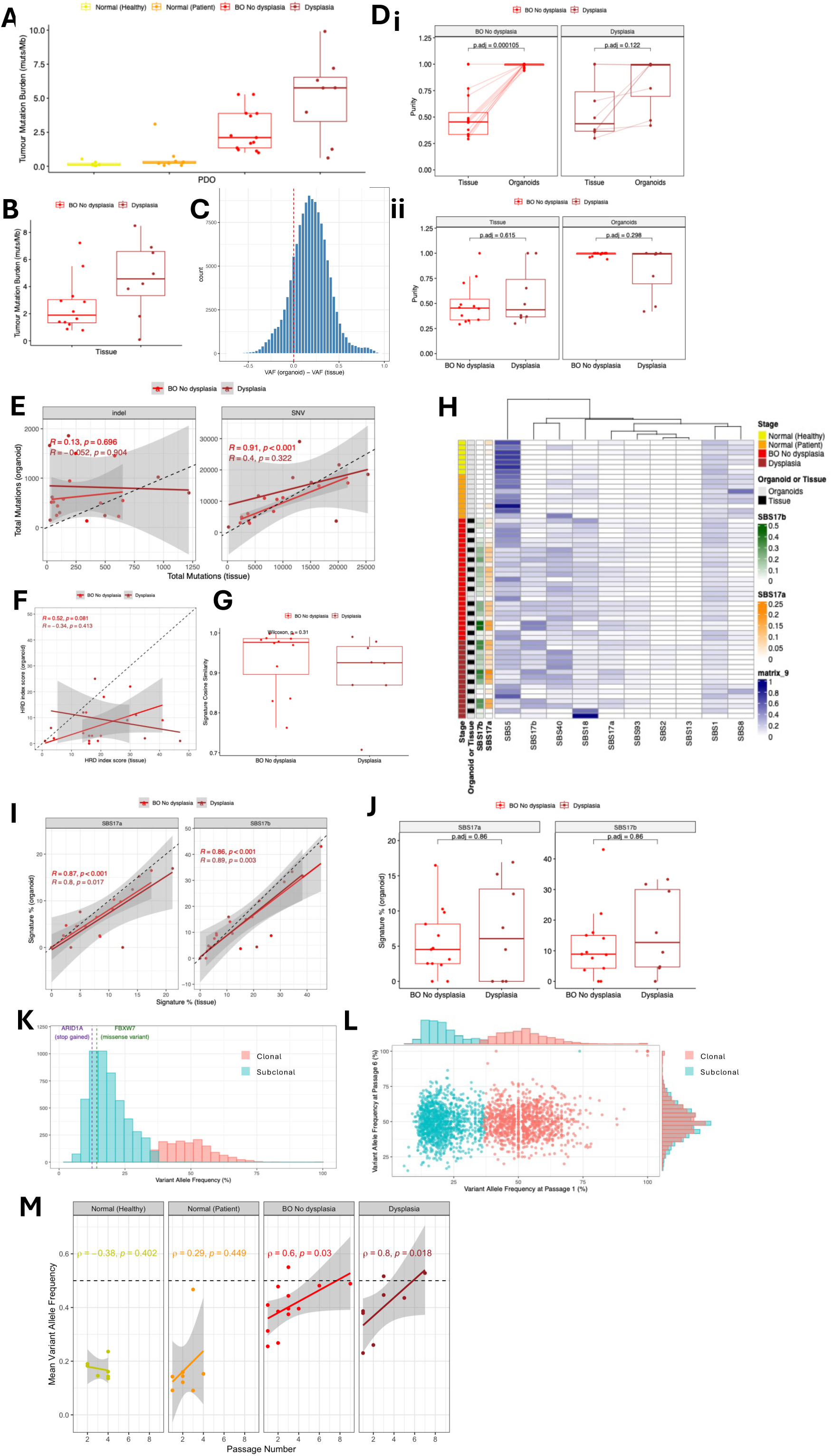
Concordance of genomic features between Barrett oesophagus PDOs and their parent tissues. Tumour mutation burden across **(A)** PDOs derived from normal gastric and BO tissue, and **(B**) tissues derived from non-dysplastic and dysplastic BO. **(C)** Differences in VAFs of shared mutations across PDO-tissue pairs, with the dotted red-line indicating mutations where VAF_organoid –_ VAF_tissue_ = 0 (i.e. where a mutation has equal VAF in a PDO and its adjacent tissue). **(D) (i)** Comparisons of tumour purity between adjacent tissues and PDOs, and **(ii)** comparisons of tumour purity between non-dysplastic and dysplastic Barrett oesophagus. Purity within PDO-tissue pairs was calculated via a pairwise Wilcoxon test, and purity across Barrett dysplastic stages was compared via a two-sample Wilcoxon test. **(E)** Comparisons of indel and SNV burdens across PDO-tissue pairs. Correlation strengths were reported as Pearson correlation coefficients. The black dotted line represents the line where (i.e. where the number of mutations is equal between a PDO and its adjacent tissue). **(F)** Comparisons of HRD index scores across PDO-tissue pairs, with Pearson correlation coefficients reported for non-dysplastic dysplastic BO derived samples. The black dotted line represents the line where *HRD_organoid_* = *HRD_tissue_* (i.e. where the HRD index scores for a PDO is equal to that of its adjacent tissue). **(G)** Comparisons of cosine similarity scores of signature profiles as reported in Figure 1F and **Supplementary Figure 1D** across non-dysplastic and dysplastic Barrett oesophagus-derived PDO-tissue pairs. Similarity scores were compared using a two-sample Wilcoxon test. **(H)** Summary of mutational signature analysis on adjacent Barrett oesophagus-derived PDOs and tissues. Samples (rows) are ordered into PDO-tissue pairs. **(I)** Correlations in SBS17a/b contributions across PDO-tissue pairs with Pearson correlation coefficients calculated separately across non-dysplastic and dysplastic samples. **(J)** Comparisons of SBS17a/b contributions across non-dysplastic and dysplastic Barrett oesophagus-derived PDOs. Contributions were compared using two-sample Wilcoxon tests. Correction for multiple testing was conducted using the Benjamini -Hochberg procedure. **(K)** VAF distributions of AHM0040, a PDO derived from a non-dysplastic Barrett oesophagus case. Dotted lines represent the VAFs of mutations in OAC-associated driver genes. **(L)** VAF distributions of shared mutations at passages 1 and 6 of AHM1373, a non-dysplastic derived PDO. Colours refer to the clonal classifications of mutations at passage 1. **(M)** Average VAF distrubtion of all normal and precancerous PDOs across their passage numbers, Spearman’s rank correlation.

**Supplementary Figure 3:**
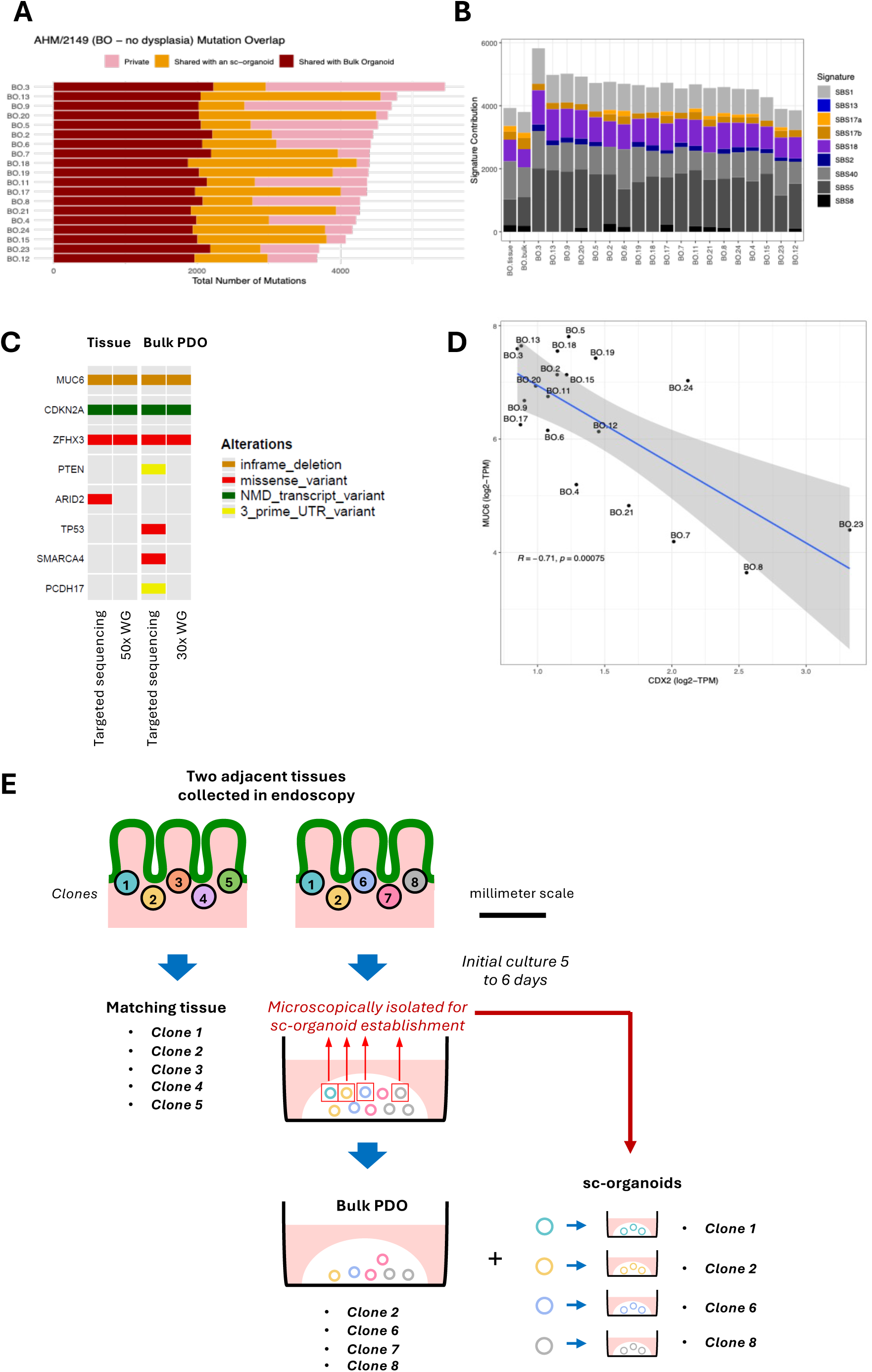
Clonal expansions of Barrett PDOs across passages and within a non-dysplastic Barrett-derived panel of single cell-derived clonal organoids. **(A)** Tumour mutation burdens of the single cell-derived clonal organoid panel derived from a non-dysplastic Barrett oesophagus PDO (AHM2149), with mutations classified as those shared with the bulk organoid from which they were derived, those not captured in the bulk organoid but present in at least one derived organoid, and private mutations. **(B)** Signature contributions across all mutations from the single cell-derived clonal organoid panel and its adjacent tissue and bulk organoid. **(C)** Comparisons of WGS versus targeted amplicon sequencing for capturing driver mutations identified in the sc-organoid panel. **(D)** Correlation between *MUCc* (gastric marker) and *CDX2* (intestinal marker). **(E)** Schematic illustrating how sc-organoids are established and their clonal relatedness to bulk PDOs and matching tissue. Note that the clonal populations present across matching tissue, bulk PDOs, and sc-organoids are not necessarily identical.

**Supplementary Figure 4:**
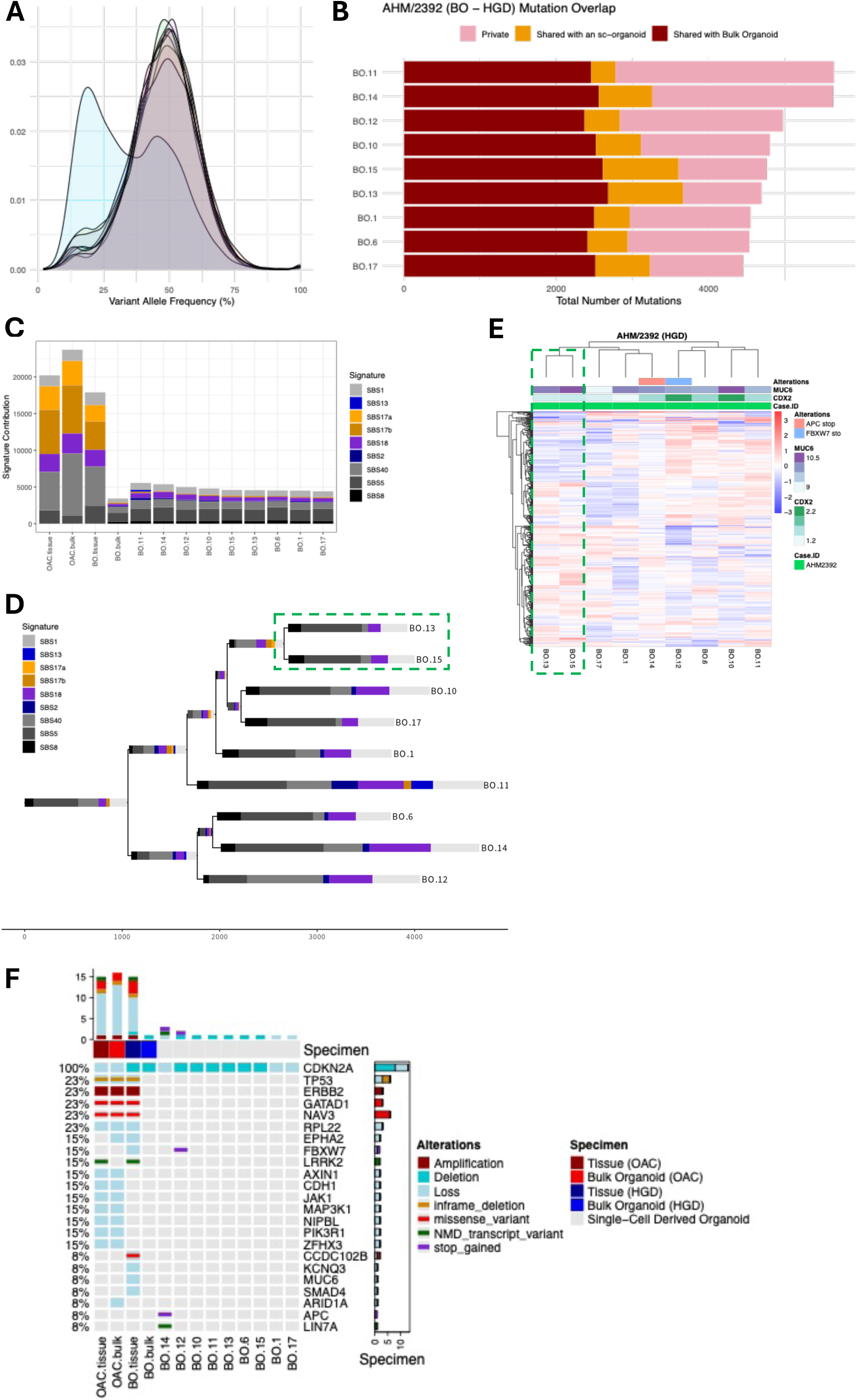
Multi-omic profiling of a single cell-derived clonal organoid panel obtained from a dysplastic Barrett oesophagus PDO. **(A)** VAF distributions across a single cell-derived clonal organoid panel (n=9) derived from a dysplastic PDO (AHM2392). Normal distributions around VAF=50% indicate clonality. **(B)** Tumour mutation burdens of the dysplastic-derived single cell-derived clonal organoid panel, with mutations classified as those shared with the bulk organoid from which they were derived, those not captured in the bulk organoid but present in at least one derived organoid, and private mutations. **(C)** Signature contributions across all mutations from the derived organoid panel, alongside adjacent tissue and bulk PDO obtained from the dysplastic and OAC tissues. **(D)** A phylogenetic profile of the PDO derived from AHM2392 defined using the derived organoid panel. Signature contributions were calculated across mutations contributing to each branch, respectively. Branch lengths are generally proportional to total amount of mutations. Signatures were not fitted for branches with fewer than 25 SNVs. **(E)** Hierarchical clustering of RNA-seq profiling for the dysplasia single cell-derived clonal organoid panel using the top 500 variable genes following variance stabilised transformation of read counts. Heatmap annotation includes genetic alterations as reported in (F) and log2-transformed TPM-normalised counts of *MUCc* (a gastric marker) and *CDX2* (an intestinal markers). The green dotted lines in **(D)** and **(E)** represent matching clones according to the phylogenetic profiling and hierarchical clustering. **(F)** Driver alterations found across the dysplasia single cell-derived clonal organoid panel as well as bulk PDO and tissue derived from dysplasia and matched OAC tissue.

**Supplementary Figure 5:**
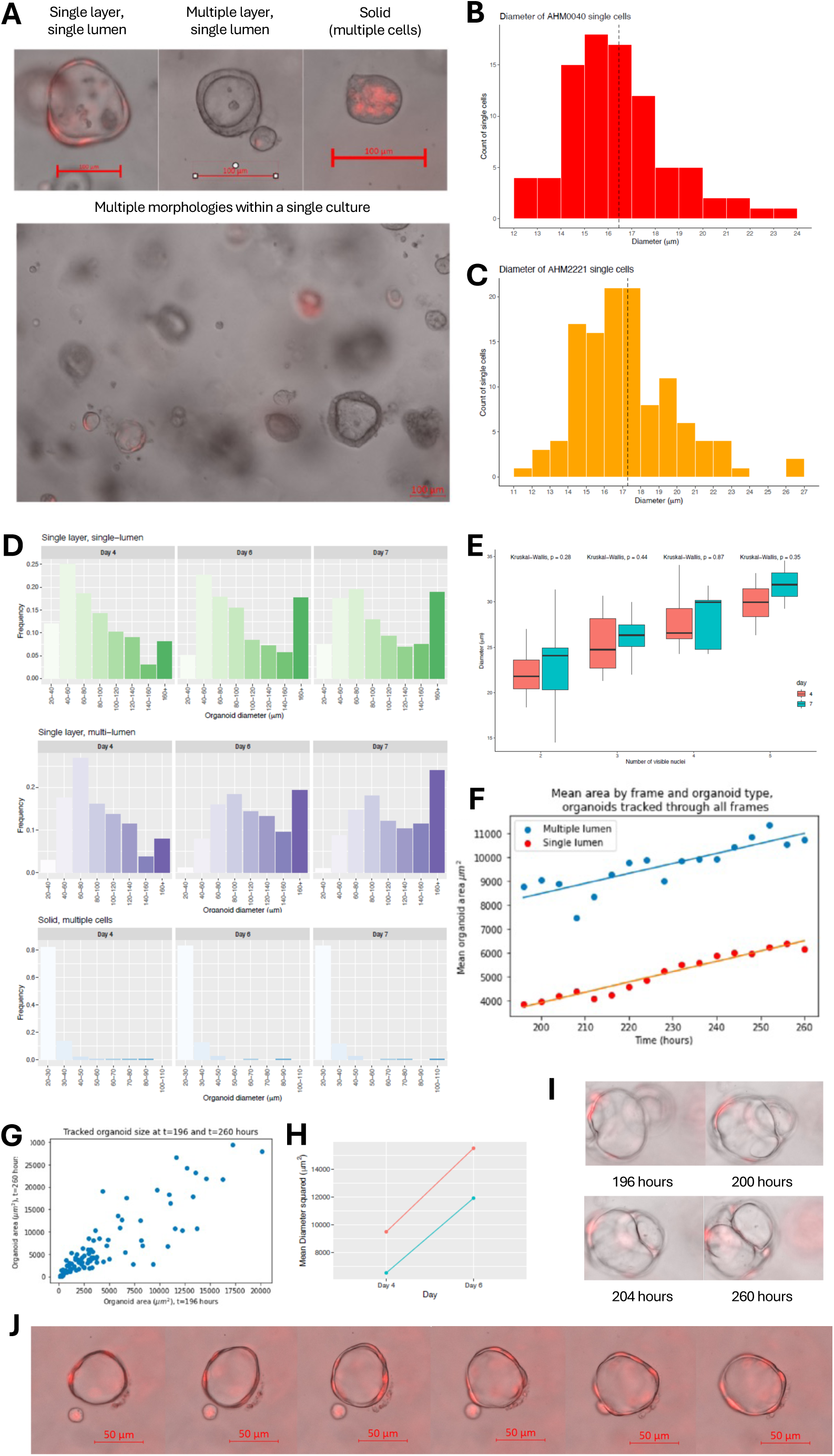
Morphological analysis of Barrett oesophagus PDOs. **(A)** Representative bright-field images illustrating the three morphology classes visible in a dysplastic Barrett oesophagus PDO (AHM2221): single layer, single lumen; multiple layer, single lumen; and solid (multiple cells, no visible lumen). Lower panel shows coexistence of multiple morphologies within a single culture. Scale bars: 100 µm (top), 200 µm (bottom). **(B and C)** Size distribution of single cells identified in the non-dysplastic (AHM0040) and dysplastic (AHM2221) Barrett oesophagus PDO cultures. Histograms show counts per 1µm; dashed vertical line marks the mean cell diameter. PDOs were counted as a diameter of 21µm or greater. **(D)** Diameter distributions of measured non-dysplastic (AHM0040) PDOs stratified by morphology class and day (within each class, lighter to darker bars denote increasing time). Each subplot shows the frequency (proportion of organoids per bin) for that class/day. **(E)** Relationship between non-dysplastic (AHM0040) PDO size and proliferative activity. Boxplots show organoid diameter versus the number of nuclei scored per organoid, colour-coded by day (as labelled). Boxes indicate interquartile range, with the median value shown as the centre line. **(F)** Mean non-dysplastic (AHM0040) PDO area over time, separated by morphology (multiple lumen, blue; single lumen, red). **(G)** Adjacent non-dysplastic (AHM0040) PDO sizes measured at 196 hours and 260 hours after seeding. Each point is one PDO; displacement from the diagonal reflects growth between time points. **(H)** Mean diameter-squared (proportional to number of cells) of single layer, single lumen organoids for non-dysplastic (AHM0040) and dysplastic (AHM2221) Barrett oesophagus organoids 4 and 6 days following seeding. **(I)** Time-lapse example of fusion in non-dysplastic (AHM0040) PDOs producing a single-layer, multi-lumen (SLML) morphology: at 196 hours the two organoids are separate but by 204 hours they have fused; the fused organoid maintains an SLML morphology through 260 hours. **(J)** Example of a non-dysplastic (AHM0040) singe-layer single-lumen PDO interacting with and likely fusing with a single cell: in the fifth frame the single cell aligns with the organoid; in the sixth frame the single cell is no longer visible, and the organoid shows a thickened fluorescent region at the point of contact.

**Supplementary Figure 6:**
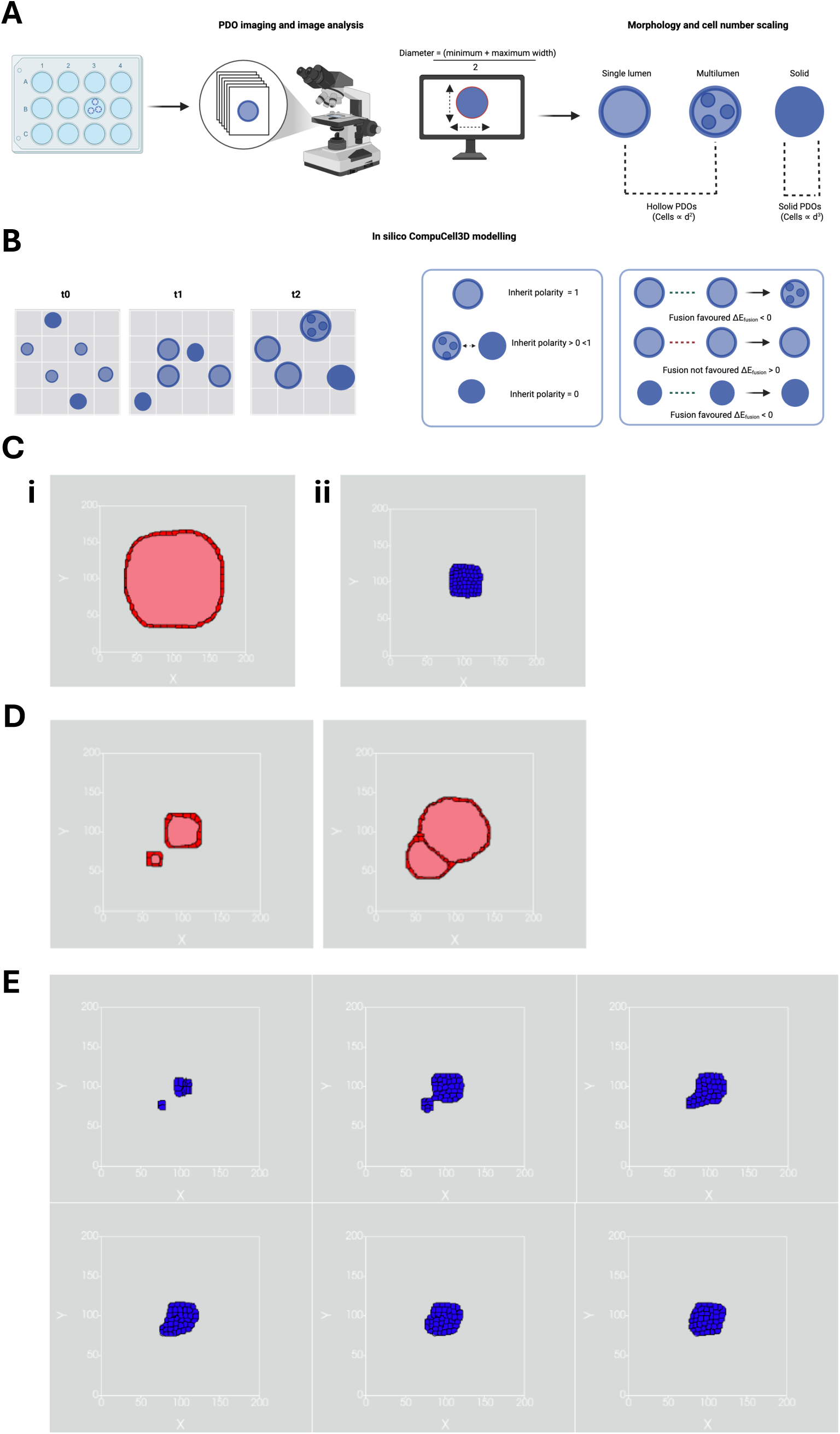
*In silico* modelling of PDO growth with variable polarity and polarity-inheritance. (A and. **B)** Overview of experimental quantification and *in silico* modelling of PDO morphology. **(A)** PDOs are imaged as z-stacks; the best-focused plane is outlined with an ellipse in Fiji and diameter is taken as the mean of minimum and maximum widths. PDOs are classified as single lumen, multi-lumen or solid; cell number is assumed to scale with diameter² for hollow organoids and diameter³ for solid PDOs **(B)** A 2D Cellular Potts model (CompuCell3D) simulates organoid cross-sections over Monte Carlo steps, illustrating growth, movement and fusion over timepoints t0, t1, and t2. Each new PDO is assigned a polarity state according to an inherited polarity parameter, IP: IP=1 yields polarised, single-lumen, single-layer PDOs; IP >0 but <1 produces a mixture of lumen-forming and solid PDOs; and IP=0 produces non-polar, solid, non-luminal PDOs. Fusion events are favoured when the change in effective adhesion energy for increasing organoid–organoid contact is negative (ΔE_fusion_ <0). **(C)** Corresponding steady-state morphologies from the model: (i) a polarised PDO forming a single lumen and single epithelial layer (IP = 1), and (ii) a non-polar PDO developing as a solid, non-luminal structure (IP = 0). **(D)** Example of simulated PDO fusion: two separate single-lumen PDOs at an early time point (left) and the resulting fused multi-lumen PDO at a later time point (right). **(E)** Time course of a representative fusion event in the model, illustrating the approach, contact and remodelling of two solid PDOs into a single composite structure over successive Monte Carlo steps.

**Supplementary Figure 7:**
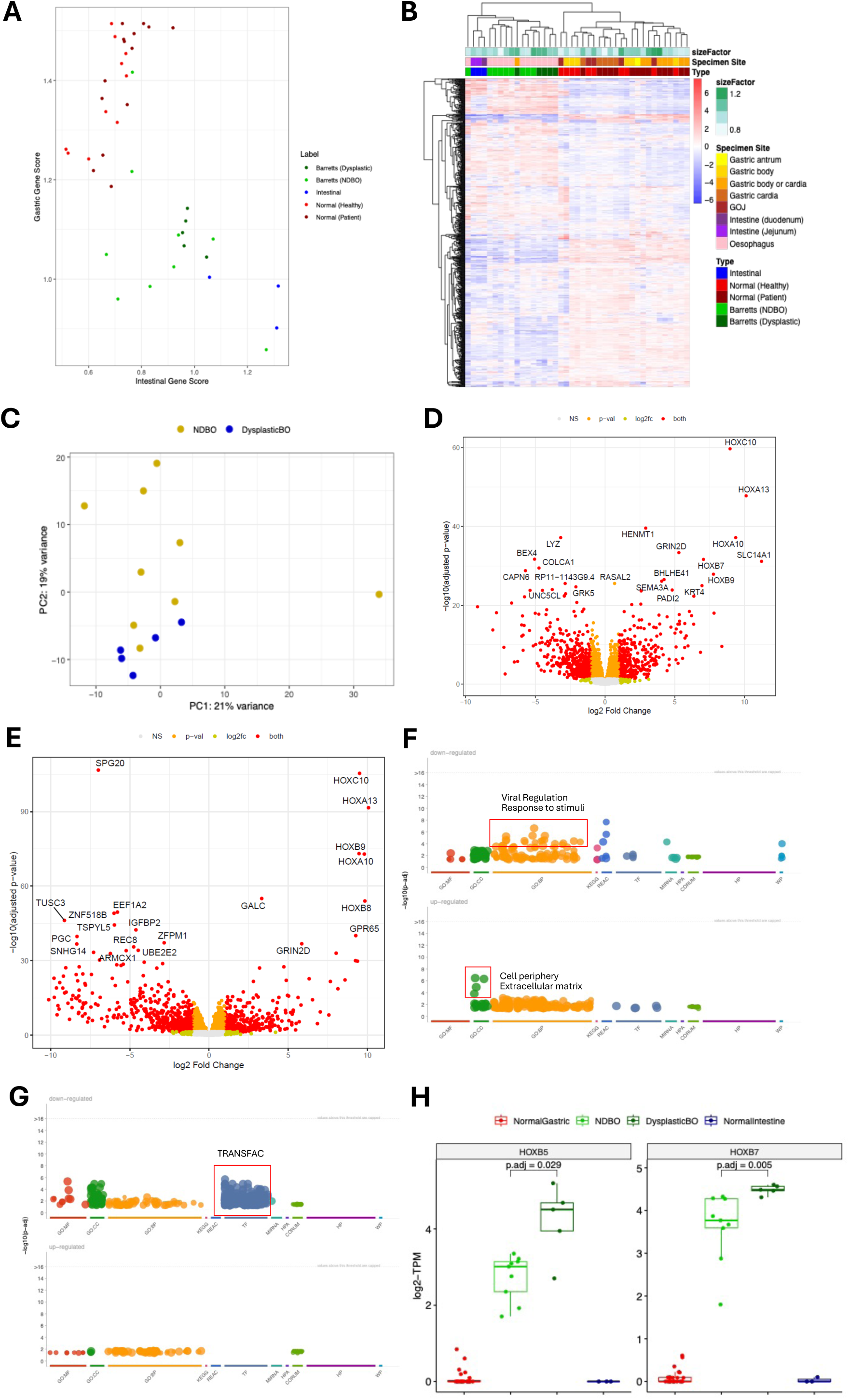
Variation in transcriptional profiles across normal- and Barrett oesophagus-derived PDOs. **(A)** Comparisons of gastric and intestinal gene panel scores across the PDO cohort, calculated via gene set variation analysis on gene panels derived from the literature. **(B)** Unsupervised hierarchical clustering of PDOs using the top 1000 variable genes. Expression values underwent variance stabilising transformation followed by subtraction of the average expression for the respective gene. **(C)** Principal component analysis of Barrett oesophagus-derived PDOs alone. **(D and E)** Volcano plots displaying the results of differential expression analysis comparing (D) non-dysplastic and (E) dysplastic Barrett oesophagus-derived PDOs with normal gastric PDOs. Genes were coloured with relation to adjusted p-values less than 0.05 and absolute log2-fold change greater than 1. The 20 genes with the lowest adjusted p-values are annotated. **(F)** Functional enrichment analysis of up- and down-regulated DEGs specific to non-dysplastic PDOs. Full results are reported in **Supplementary Table 7**. **(G)** Functional enrichment analysis of up- and down-regulated DEGs specific to dysplastic PDOs. Full results are reported in **Supplementary Table 8**. **(H)** Comparisons of *HOXB5* and *HOXB7* gene expression across the PDO cohort. TPM-values were reported following log2-transformation. Gene expression was compared between non-dysplastic and dysplastic PDOs using a two-sample Wilcoxon test, adjusted for multiple testing using the Benjamini-Hochberg procedure.

**Supplementary Figure 8:**
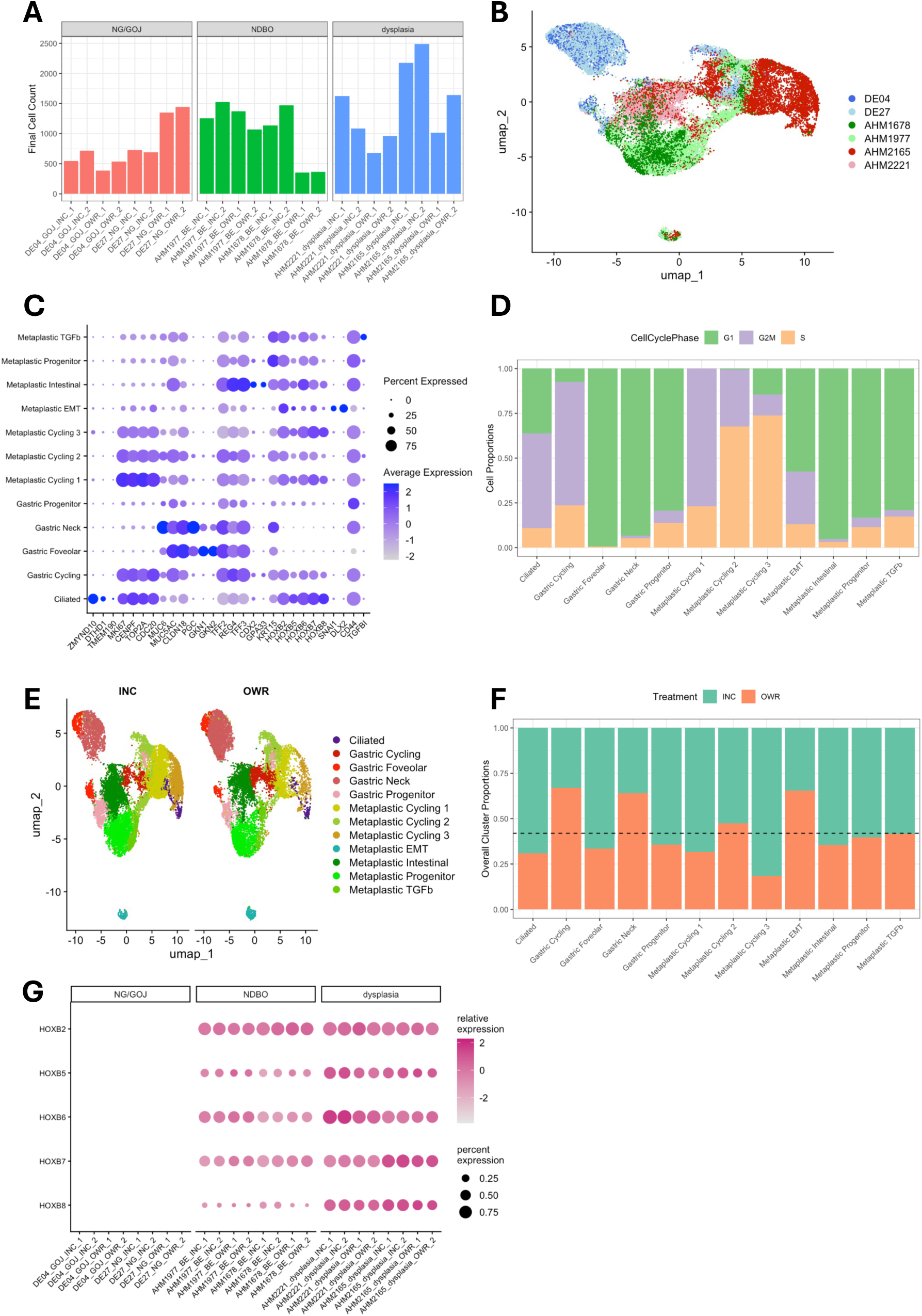
Single-cell RNA-seq analysis of normal gastric, BO-, and dysplasia-derived PDOs. **(A)** Final counts of sequenced cells across 24 PDOs included in analysis. **(B)** UMAP dimensionality reduction of all cells coloured by sample. **(C)** Gene markers used for defining phenotypes of each cell cluster. **(D)** Cell cycle phase classification of cells across each cluster. **(E)** UMAP dimensionality reduction of cells split by applied organoid media. **(F)** Comparison of cell counts from PDOs treated with INC and OWR culture media, separated by cell cluster. **(G)** HOXB genes expression across 24 PDO samples.

**Supplementary Figure 9:**
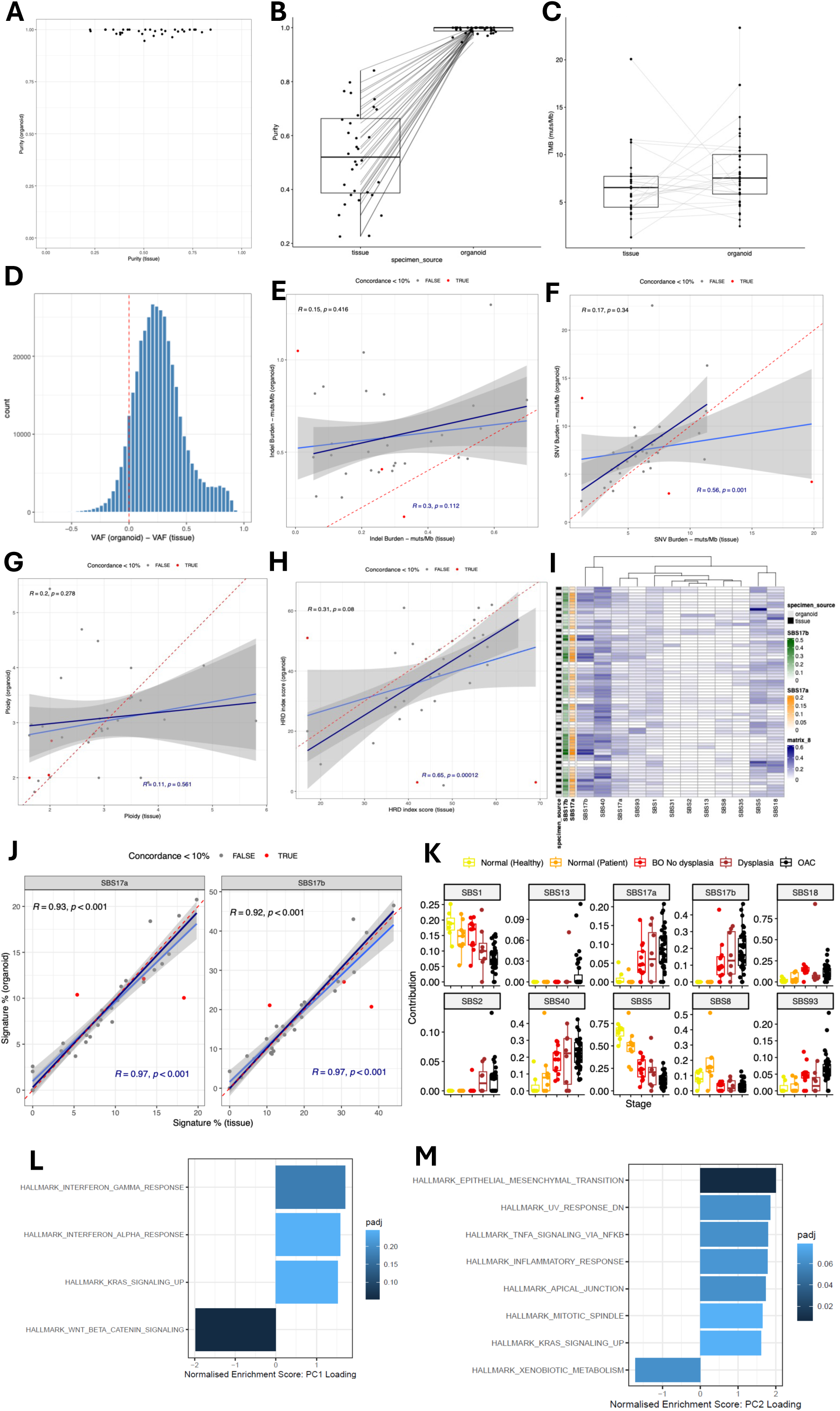
Genomic features of OAC PDOs and their relationship to parent tissue. (A and. **B)** Correlation between tumour purity for PDOs and their adjacent tissues. Purity between PDO-tissue pairs was calculated via a pairwise Wilxocon test. **(C)** Tumour mutational burden in PDOs and their adjacent tissues. **(D)** Differences in VAFs of shared mutations across PDO-tissue pairs, with the dotted red-line indicating mutations where VAF_organoid –_ VAF_tissue_ = 0 (i.e. where a mutation has equal VAF in a PDO and its adjacent tissue). **(E)** Comparison of insertion-deletion (indel) mutation burden between PDOs and their adjacent tissues. **(F)** Comparison of SNV burden between PDOs and their adjacent tissues. **(G)** Comparison of ploidy in PDOs and their adjacent tissues. **(H)** Comparison of HRD index score in PDOs and their adjacent tissues. **E-H)** Red dots indicate PDO-tissue pairs with mutational concordance less than 10% and gray dots indicate PDO-tissue pairs with mutational concordance more than 10%, as reported in Figure 5C. Red dotted lines indicate equivalent values (i.e. y=x). Solid blue lines and shaded grey areas indicate linear regression lines of best fit alongside 95% confidence bands. Solid navy lines indicate linear regression lines following the removal of PDO-tissue pairs with mutational concordance less than 10%. Comparisons were made using Pearson correlation coefficients with R- and p-values shown. Values shown in black in the top left were calculated across all PDO-tissue pairs, and values shown in navy in the bottom right were calculated following the removal of PDO-tissue pairs with mutational concordance less than 10%. **(I)** Summary of mutational signature analysis on adjacent OAC PDOs and their parent tissues. Samples (rows) are ordered into matched PDO-tissue pairs. **(J)** Correlations in SBS17a (left) and SBS17b (right) contributions across PDO-tissue pairs with Pearson correlation coefficients calculated separately for SBS17a and SBS17b. Annotations are the same as in (E to H). **(K)** Contributions of mutational signatures across both the gastric/BO and OAC PDO cohorts. **(L and M)** Gene set enrichment analysis of the loadings for (L) PC1 and (M) PC2 obtained from principal component analysis of RNA-seq data from the OAC PDO cohort. Analysis was based on the top 1000 variable genes across the cohort.

**Supplementary Figure 10:**
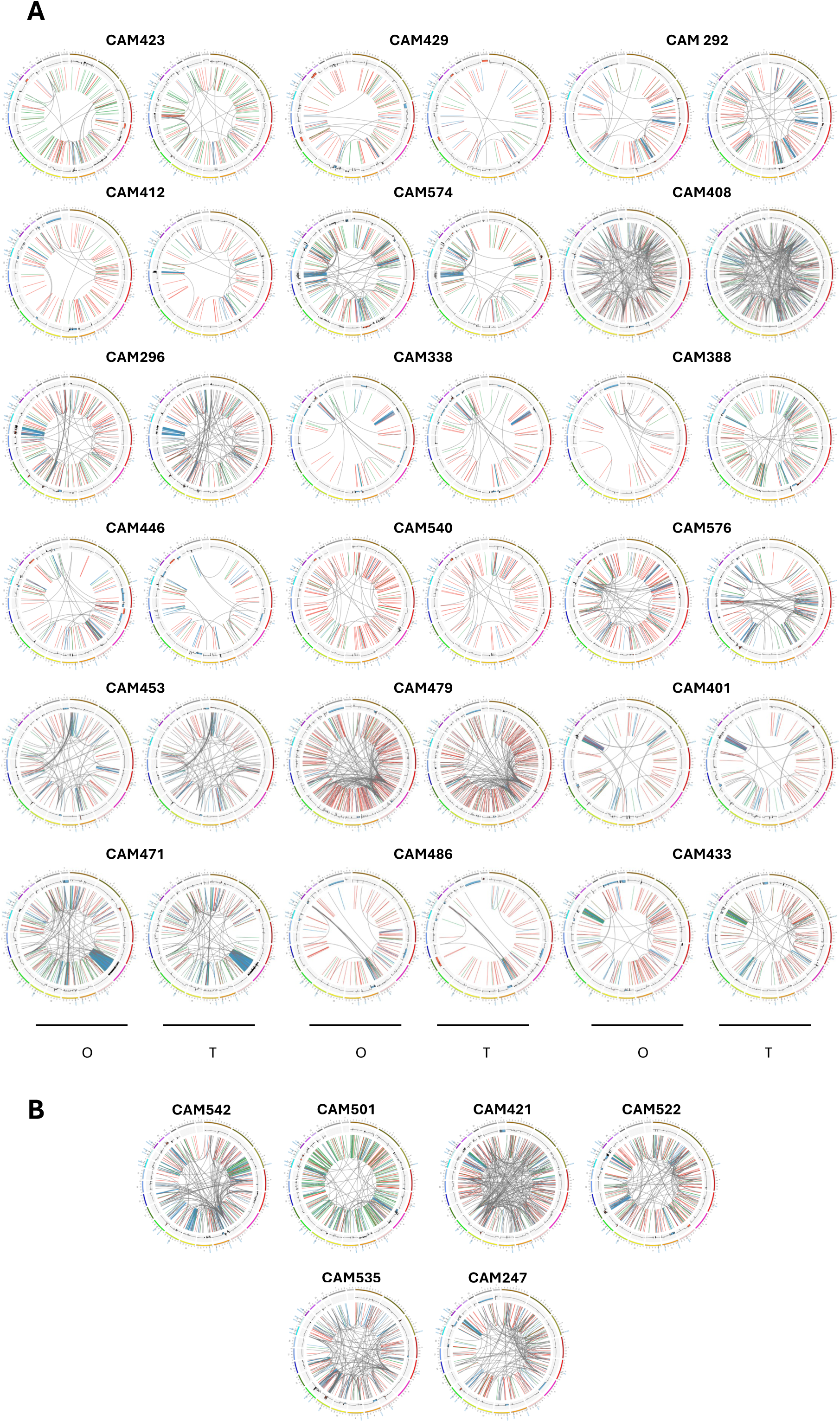
Circos plots demonstrating genomic similarities between each OAC PDO and its parent tissue. Each plot demonstrates overall genomic landscape of the organoid (O) and adjacent tumour tissue (T). The outer circle represents the labelled chromosomes with labelled driver gene positions. The middle circle depicts copy number profile where orange shows gains and blue shows deletions. The inner most circle shows inter and intra chromosomal structural variations with grey, red, blue and green lines representing translocations, deletions, inversions and duplications, respectfully.

**Supplementary Figure 11:**
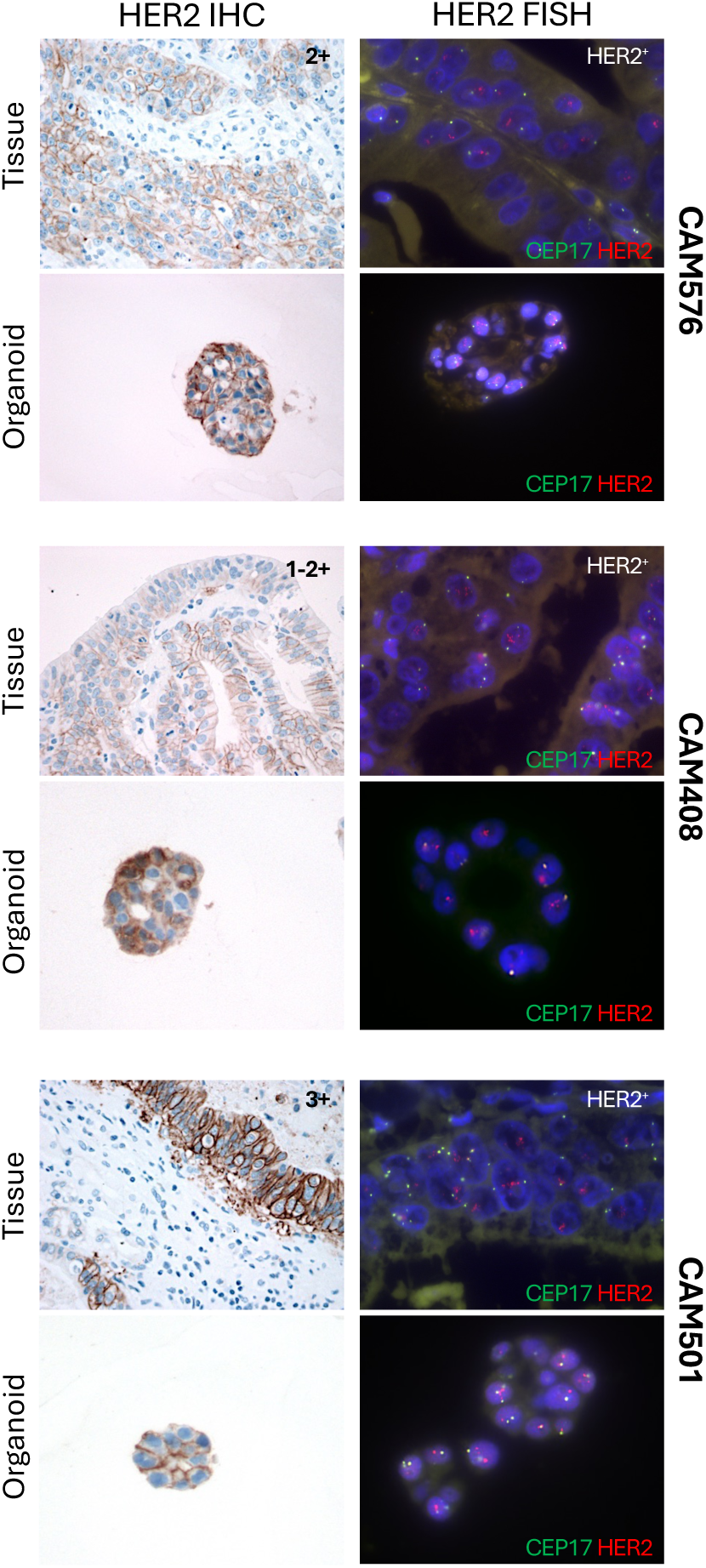
Immunohistochemistry and fluorescence *in situ* hybridisation of *HER2*. IHC and FISH analysis of suspected HER2amplification in PDOs and matched tissue. IHC scores are shown for each tissue image based on current HER2 OAC diagnostic guidelines (0 negative, 1-2+ equivocal, 3+ positive). FISH probes and colours are indicated in the bottom left of each FISH image with chromosome 17 (CEN17q) probe shown as green and the HER2probe as red. Positive HER2 amplification determined by FISH is denoted as HER2+ in each image. All organoids and their derivation tissue are grouped together, and case IDs are labelled on the right.

**Supplementary Figure 12:**
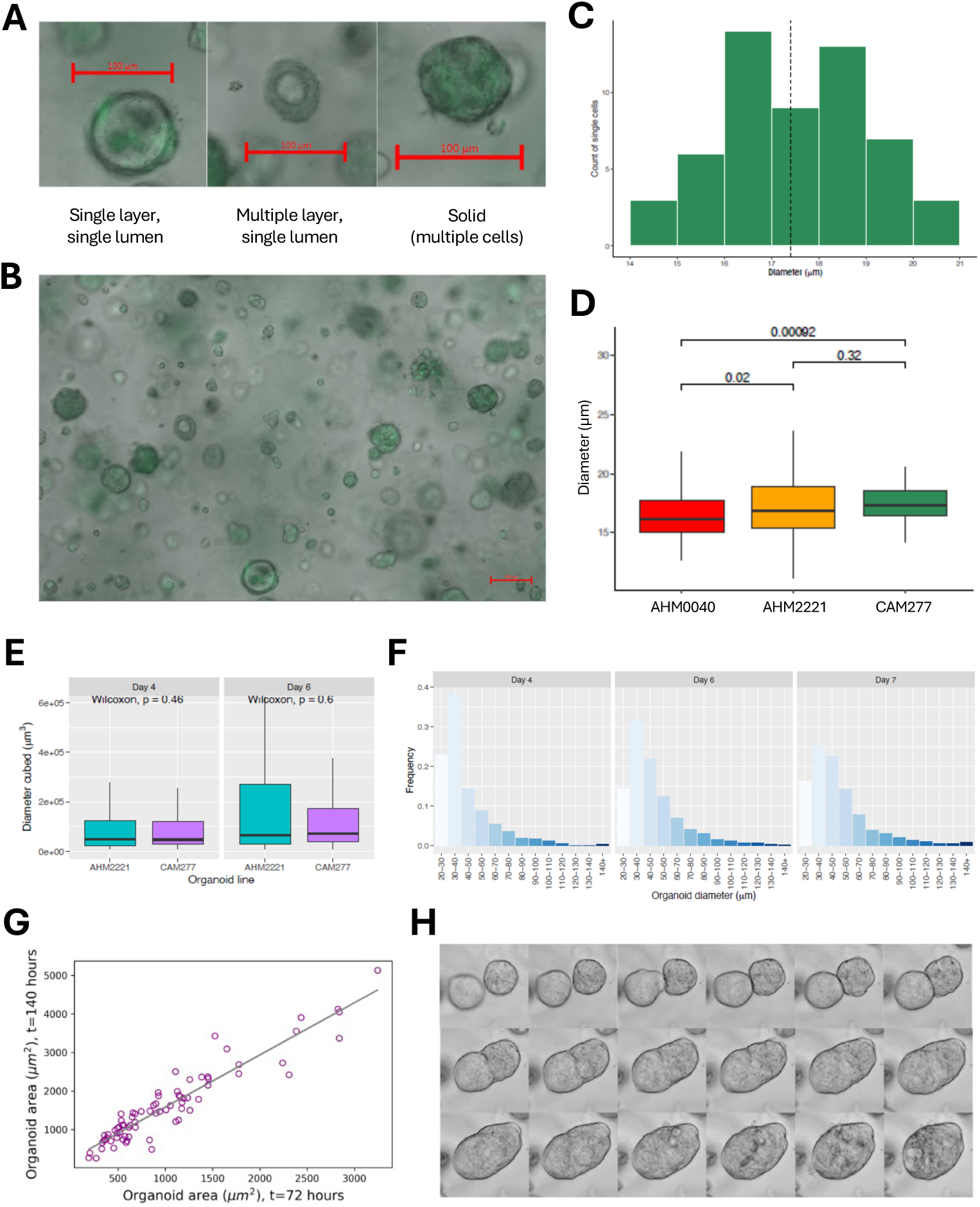
Morphological analysis of OAC PDOs. **(A)** Representative bright-field images illustrating the three morphology classes visible in an OAC PDO (CAM277): single layer, single lumen; multiple layer, single lumen; solid. Image taken at day 4. Scale bar shows 100µm. **(B)** Coexistence of multiple morphologies at day 7 of a single CAM277 culture. Scale bar shows 100µm. **(C)** Size distribution of single cells identified in a single CAM277 OAC PDO culture. Histogram shows counts per 1µm interval; dashed vertical line marks mean cell diameter. PDOs were counted as a diameter of 21µm or greater. **(D)** Size of OAC PDO (CAM277) single cells compared with Barrett oesophagus non-dysplastic (AHM0040) and dysplastic (AHM2221) PDOs. Statistical comparisons made using the Kruskal-Wallis test. **(E)** Distribution of CAM277 PDO size at days 4, 6 and 7 after seeding. **(F)** Correlation between growth at 72 hours and growth at 140 hours for 76 individual CAM277 organoids. **(G)** Representative brightfield image demonstrating fusion of two solid CAM277 organoids between 72 and 140 hours of seeding (images taken at four hourly intervals from top left through to bottom right).

**Supplementary Figure 13:**
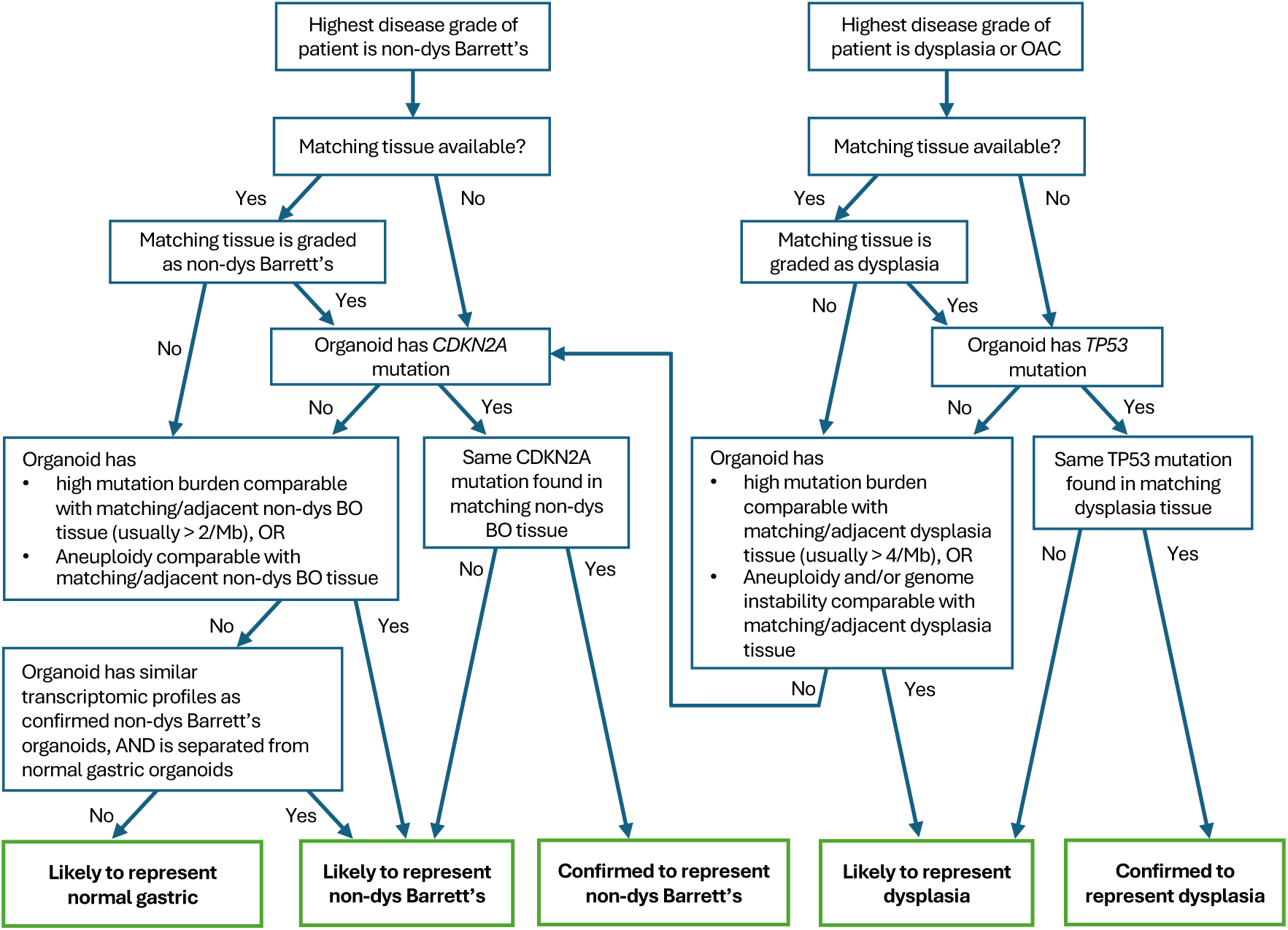
Decision framework for classifying precancerous organoids. Decision tree showing how patient-derived organoids are classified by integrating patient disease grade, matching tissue, *TP53* status, mutation burden, and genomic instability. The framework emphasizes tissue-anchored, multi-modal interpretation rather than single-feature classification, and mainly to distinguish normal gastric and precancerous organoids in BO.

## LIST OF SUPPLEMENTARY TABLES

**Supplementary Table 1:** Demographic and clinicopathological characteristics of donors for PDOs obtained from non-dysplastic and dysplastic Barrett oesophagus. - missing data. F, female; HGD, high grade dysplasia; IM, intestinal metaplasia; M, male; LGD, low grade dysplasia; NDBO, non-dysplastic Barrett oesophagus; OAC, oesophageal adenocarcinoma. *, this patient was thought to have dysplasia, but histopathology confirmed the as indefinite for dysplasia (IND).

| Patient ID | Age | Sex | Ethnicity | Organoid derived from | Highest grade at the time of tissue collection | Adjacent to OAC | Experiments |
|---|---|---|---|---|---|---|---|
| AHM0040 | 69 | M | White British | NDBO | NDBO | - | WGS, RNA-seq, live imaging |
| AHM0931 | 73 | M | White British | NDBO | NDBO | - | WGS |
| AHM1345 | 71 | F | White British | NDBO | NDBO | - | WGS, RNA-seq |
| AHM1369 | 80 | M | White British | NDBO | NDBO | - | WGS, RNA-seq |
| AHM1678 | 79 | M | White British | NDBO | NDBO | - | WGS, RNA-seq, scRNA-seq |
| AHM1977 | 68 | F | White British | NDBO | NDBO | - | WGS, RNA-seq, scRNA-seq |
| AHM2172 | 73 | F | White British | NDBO | NDBO | - | WGS, RNA-seq |
| AHM2575 | 79 | F | White British | NDBO | NDBO | - | WGS |
| AHN0070 | 83 | M | White British | NDBO | NDBO | - | WGS, RNA-seq |
| AHM1373 | 74 | M | White British | NDBO | NDBO | - | WGS, RNA-seq |
| AHM2149 | 67 | M | White British | NDBO | NDBO | - | WGS, RNA-seq |
| CAM673 | 66 | M | White British | NDBO | OAC | - | WGS, RNA-seq |
| AHM0593 | 74 | F | White British | HGD | HGD | - | WGS, RNA-seq |
| CAM478 | 61 | F | White British | LGD | OAC | Yes | WGS, RNA-seq |
| AHM2165 | 77 | M | White British | HGD_1 | OAC | Yes | WGS, RNA-seq, scRNA-seq |
| AHM2165 | 77 | M | White British | HGD_2 | OAC | - | WGS, RNA-seq |
| CAM603 | 74 | M | White British | HGD | OAC | Yes | WGS, RNA-seq |
| AHM2221 | 74 | F | White British | HGD | HGD | - | WGS, RNA-seq, scRNA-seq, live imaging |
| AHM2392 | 87 | F | White British | HGD | OAC | Yes | WGS, RNA-seq |
| AHM2688* | 73 | M | White British | IND | IND | - | WGS |
| AHM2688* | 73 | M | White British | NDBO | IND | - | WGS |

**Supplementary Table 2:** A summary of demographic and clinicopathological characteristics of normalgastric and intestinal organoids. GOJ: gastroesophageal junction. The jejunum and duodenum organoids were only used as intestinal reference in RNA-seqexperimentsin Figure 4.

| Individual ID | Age | Sex | Organoid derived from | Condition/Procedure | Experiments |
|---|---|---|---|---|---|
| DE04 | 63 | Male | GOJ | Gastrointestinal disease-free organ donor | RNA-seq, WGS, scRNA-seq |
| DE21 | 74 | F | GOJ | Gastrointestinal disease-free organ donor | RNA-seq |
| DE21 | 74 | F | Gastric body | Gastrointestinal disease-free organ donor | RNA-seq |
| DE22 | 68 | F | Gastric body | Gastrointestinal disease-free organ donor | RNA-seq |
| DE26 | 61 | Female | GOJ | Gastrointestinal disease-free organ donor | RNA-seq, WGS |
| DE26 | 61 | Female | Gastric body | Gastrointestinal disease-free organ donor | RNA-seq, WGS |
| DE27 | 35 | Male | GOJ | Gastrointestinal disease-free organ donor | RNA-seq, WGS |
| DE27 | 35 | Male | Gastric body | Gastrointestinal disease-free organ donor | RNA-seq, WGS, scRNA-seq |
| DE28 | 37 | Male | GOJ | Gastrointestinal disease-free organ donor | RNA-seq, WGS |
| DE28 | 37 | Male | Gastric body | Gastrointestinal disease-free organ donor | RNA-seq, WGS |
| AHM2221 | 74 | F | Gastric cardia | Endoscopy | RNA-seq, WGS |
| AHM0040 | 69 | M | Gastric cardia | Endoscopy | RNA-seq, WGS |
| AHM1373 | 74 | M | Gastric cardia | Endoscopy | RNA-seq, WGS |
| AHM1678 | 79 | M | Gastric cardia | Endoscopy | RNA-seq, WGS |
| AHM2165 | 77 | M | Gastric cardia | Endoscopy | RNA-seq, WGS |
| AHM2331 | 76 | M | Gastric cardia | Endoscopy | RNA-seq, WGS |
| AHM0593 | 72 | F | Gastric cardia | Endoscopy | WGS |
| AHM1373 | 74 | M | Gastric cardia | Endoscopy | WGS |
| CAM0796 | 63 | F | Gastric antrum | Endoscopy | WGS |
| CAM433 | 66 | M | Gastric body | Oesophagectomy | RNA-seq |
| CAM463 | 68 | F | Gastric body | Oesophagectomy | RNA-seq |
| CAM478 | 61 | F | Gastric body | Oesophagectomy | RNA-seq, WGS |
| CAM749 | - | - | Gastric body | Oesophagectomy | RNA-seq |
| CAM549 | - | - | Gastric body | Oesophagectomy | RNA-seq |
| CAM581 | - | - | Gastric body | Oesophagectomy | RNA-seq |
| CAM584 | - | - | Gastric body | Oesophagectomy | RNA-seq |
| MS050 | 63 | Male | Jejunum | Gastrectomy | RNA-seq |
| HGH046 | 67 | Female | Jejunum | Gastrectomy | RNA-seq |
| TB16.0980 | 42 | Male | Duodenum | Pancreaticoduodenectomy | RNA-seq |

**Supplementary Table 3:** Morphological characteristics of a non-dysplastic Barrett oesophagus PDO (AHM0040) at four, six and seven days following seeding.

| **Day** | **Measure** | **Single layer, single lumen** | **Single layer, multi-lumen** | **Multi-layer, with lumen** | **Solid, multiple cells** |
|---|---|---|---|---|---|
| **4** | Number of organoids (%) | 769 (53%) | 343 (23%) | 139 (9%) | 213 (15%) |
|  | Max diameter (µm) | 378 | 380 | 160 | 84.0 |
|  | Mean diameter (µm) | 85.2 | 94.7 | 46.3 | 27.1 |
|  | Mean circularity | 1.11 | 1.12 | 1.08 | 1.06 |
| **6** | Number of organoids (%) | 720 (58%) | 326 (26%) | 68 (6%) | 119 (10%) |
|  | Max diameter (µm) | 513 | 397 | 213 | 83.3 |
|  | Mean diameter (µm) | 106 | 121 | 63.1 | 27.0 |
|  | Mean circularity | 1.12 | 1.15 | 1.08 | 1.06 |
| **7** | Number of organoids (%) | 776 (61%) | 322 (25%) | 60 (5%) | 117 (9%) |
|  | Max diameter (µm) | 376 | 421 | 143 | 104 |
|  | Mean diameter (µm) | 108 | 129 | 64.3 | 27.3 |
|  | Mean circularity | 1.13 | 1.15 | 1.10 | 1.07 |

**Supplementary Table 4:** Morphological characteristics of a dysplastic Barrett oesophagus PDO (AHM2221) at four, six and seven days following seeding.

| **Day** | **Measure** | **Single layer, single lumen** | **Multi-layer, with lumen** | **Solid, multiple cells** |
|---|---|---|---|---|
| **4** | Number of organoids | 153 (29%) | 142 (27%) | 245 (45%) |
|  | Max diameter (µm) | 202 | 91.9 | 135 |
|  | Mean diameter (µm) | 72.8 | 52.2 | 43 |
|  | Mean circularity | 1.08 | 1.07 | 1.07 |
| **6** | # organoids (%) | 241 (34%) | 212 (30%) | 255 (36%) |
|  | Max diameter (µm) | 444 | 250 | 187 |
|  | Mean diameter (µm) | 93.5 | 64.2 | 52.7 |
|  | Mean circularity | 1.12 | 1.08 | 1.07 |

**Supplementary Table 5:** Summary of *in silico* and *in vitro* observations for the growth patterns and morphology of non-dysplastic (AHM0040) and dysplastic (AHM2221) Barrett oesophagus PDOs. BO, Barrett oesophagus; EP, establish polarity (i.e. whether cells are polar); IP, inherit polarity (i.e. the likelihood of a cell inheriting polarity); NBDO, non-dysplastic Barrett oesophagus; OAC, oesophageal adenocarcinoma; SLSL, single layer, single lumen.

|  | **Single layer, single lumen** | **Single layer, multi-lumen** | **Multi-layer, with lumen** | **Solid** |
|---|---|---|---|---|
| ***In vitro* observations (>1% organoids)** | BO lines only (AHM0040, AHM2221) | NBDO only (AHM0040) | BO lines only (AHM0040, AHM2221) | BO and OAC (AHM0040, AHM2221, CAM277) |
| ***In Silico* observations** | Single organoids | Fused SLSL organoids | Single organoids | Single and fused organoids |
| **Parameters** | IP = 1<br>EP = True | As for SLSL | IP<1, IP>0<br>EP = True | IP 0 or<br>EP False |
| **Parameter interpretation** | Cells all polar | Cells all polar | Some cells with disrupted polarity | Cells all with disrupted polarity |

< Supplementary_Tables_6_7_8_14.xlsx>

**Supplementary Table 6:** a summary of enriched processes and pathways relating to genes that are differentially expressed only non-dysplastic and dysplastic Barrett oesophagus versus gastric patient derived organoids. BP: biological process; GO: gene ontology; MF: molecular function; TF: transcription factor; WP: Wiki pathway.

< Supplementary_Tables_6_7_8_14.xlsx>

**Supplementary table 7:** a summary of enriched processes and pathways relating to genes that are differentially expressed only in non-dysplastic Barrett oesophagus versus gastric patient derived organoids. BP: biological process; GO: gene ontology; MF: molecular function; TF: transcription factor; WP: Wiki pathway.

< Supplementary_Tables_6_7_8_14.xlsx>

**Supplementary table 8:** a summary of enriched processes and pathways relating to genes that are differentially expressed only in dysplastic versus gastric patient derived organoids. BP: biological process; GO: gene ontology; MF: molecular function; TF: transcription factor; WP: Wiki pathway.

**Supplementary Table 9:** Morphological characteristics of an OAC PDO (CAM277) at four, six and seven days following seeding.

| **Solid organoids** | **Day 4** | **Day 6** | **Day 7** |
|---|---|---|---|
| Number of organoids | 1936 | 2013 | 1918 |
| Mean diameter (µm) | 43.2 | 47.9 | 49.2 |
| Maximum diameter (µm) | 183 | 262 | 213 |
| Mean circularity | 1.07 | 1.07 | 1.08 |
| Proportion of solid organoids | 99% | 99% | 99% |

**Supplementary Table 10:** OAC PDO sensitivity to cytotoxic chemotherapy. AUC: area under the curve (an integral value of viability following exposure to defined concentrations of each drug); E_max_: the maximum effect; IC_50_: the concentration of each drug required to reduce viability to 50%; 5FU: 5-fluorouracil.

|  | **5FU+ Leucovorin** |  |  | **Oxaliplatin** |  |  | **Docetaxel** |  |  |
|---|---|---|---|---|---|---|---|---|---|
|  | **IC50** | **AUC** | **Emax** | **IC50** | **AUC** | **Emax** | **IC50** | **AUC** | **Emax** |
| **CAM574** | 15.66 | 194.73 | 93.05 | 20.19 | 250.85 | 91.64 | 1.68 | 154.30 | 77.71 |
| **CAM277** | 8.31 | 191.57 | 93.94 | 7.73 | 169.87 | 95.82 | 2.10 | 177.37 | 78.18 |
| **CAM408** | 51.33 | 287.0 | 95.65 | 60.71 | 253.4<sup>^</sup> | 62.89<sup>^</sup> | 3.04 | 161.75 | 72.79 |
| **CAM117** | 14.65 | 210.25 | 88.51 | 96.78 | 246.25 | 76.87 | 2.57 | 140.35 | 93.53 |
| **CAM146** | 25.83 | 222.4 | 86.6 | 92.66<sup>*</sup> | 257.0<sup>*</sup> | 74.15<sup>*</sup> | 1.64 | 214.60 | 50.04 |

\* n=1 biological replicate (three technical replicates)
^ AUC and Emax were normalized to a top concentration of 100 µM. Only one biological replicate (three technical replicates) was dosed to 200 µM

**Supplementary Table 11:**
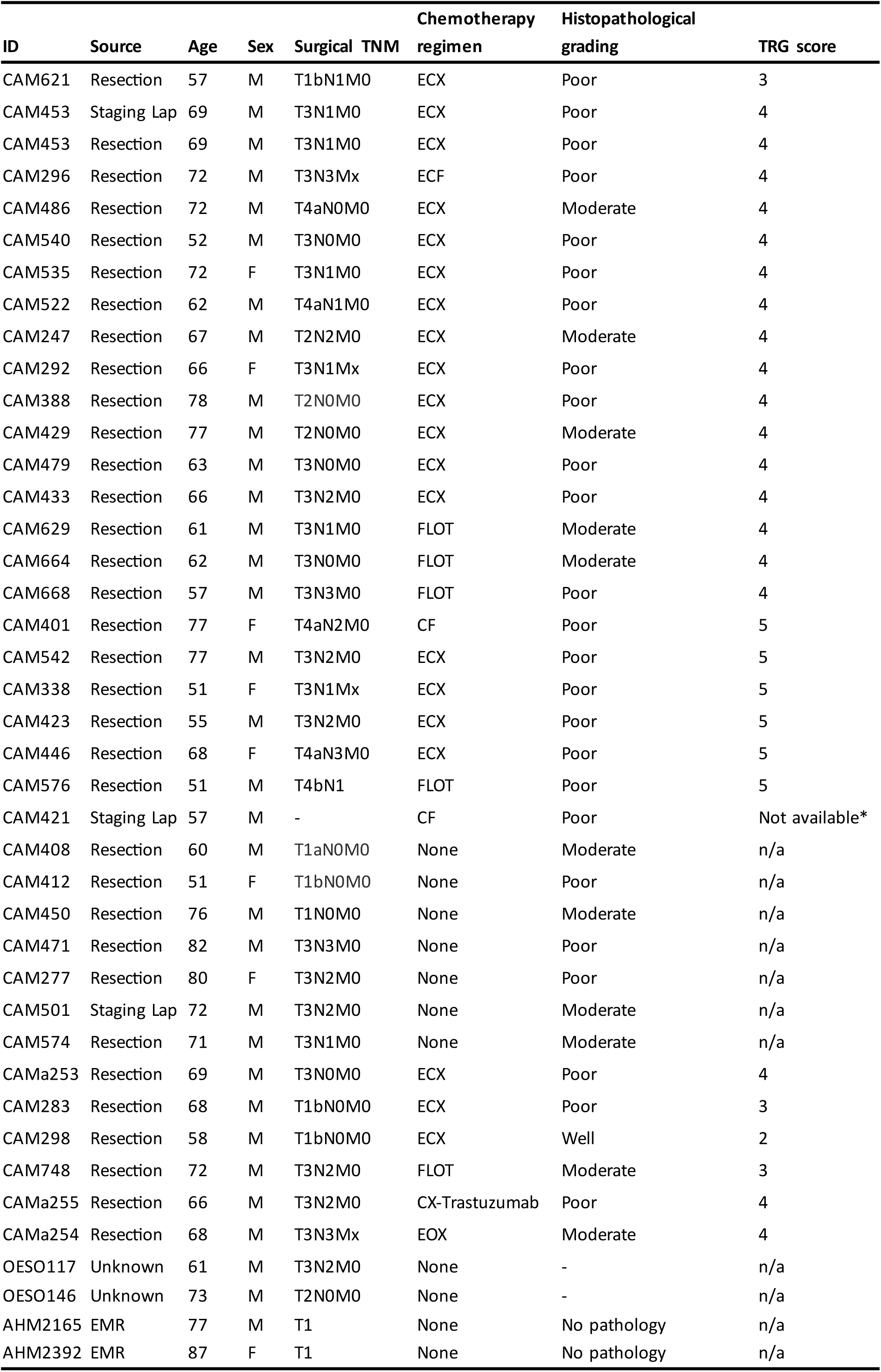
Demographic and clinicopathological characteristics of organoids derived from patients with oesophageal adenocarcinoma. All patients were White British. *TRG not available but sample noted demonstrate therapy-related tumour regression. - missing data. ECX, epirubicin, cisplatin, capecitabine; CF, cisplatin, 5-fluorouracil; FLOT: 5- fluorouracil/leucovorin, oxaliplatin, docetaxel; Lap laparoscopy; TRG, tumour regression grade.

**Supplementary Table 12:** OAC PDO sensitivity to x-ray irradiation. Organoid forming efficiency shown for each PDO following exposure to 0-8Gy x-ray irradiation in a single fraction. AUC, median area under the curve (integral of 0-8Gy organoid forming efficiency values).

|  | **CAM277** | **CAM401** | **CAM408** | **CAM412** | **CAM429** | **CAM501** | **CAM535** | **CAM574** |
|---|---|---|---|---|---|---|---|---|
| **0 Gy** | 1 | 1 | 1 | 1 | 1 | 1 | 1 | 1 |
| **2 Gy** | 0.86 | 0.88 | 1.03 | 0.77 | 0.71 | 0.42 | 0.58 | 0.62 |
| **4 Gy** | 0.80 | 0.62 | 0.99 | 0.56 | 0.66 | 0.54 | 0.57 | 0.78 |
| **6 Gy** | 0.74 | 0.77 | 1.10 | 0.52 | 0.53 | 0.36 | 0.59 | 0.70 |
| **8 Gy** | 0.68 | 0.75 | 0.73 | 0.62 | 0.60 | 0.24 | 0.47 | 0.55 |
| **AUC** | 0.83 | 0.79 | 0.98 | 0.66 | 0.67 | 0.46 | 0.63 | 0.70 |

**Supplementary Table 13:**
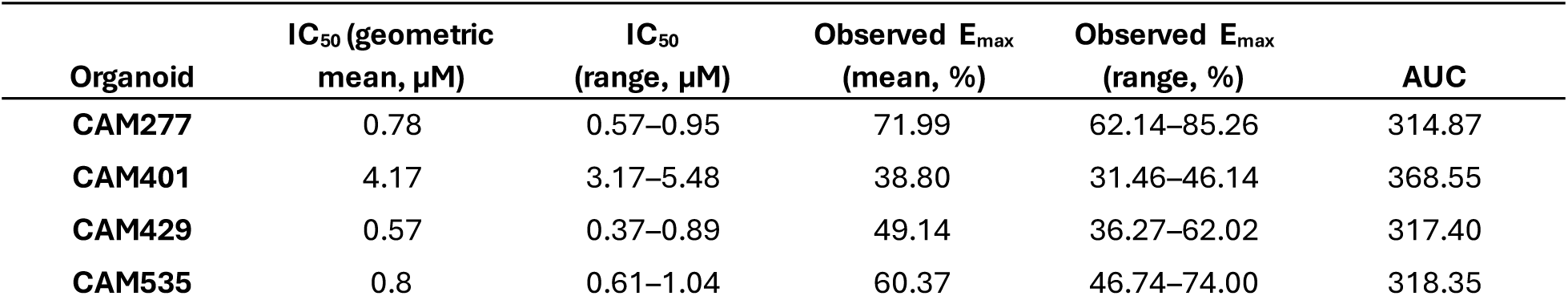
OAC PDO sensitivity to the CDK4/6 inhibitor abemaciclib. AUC: area under the curve (an integral value of viability following exposure to defined concentrations of each drug); E_max_: the maximum effect; IC_50_: the concentration of each drug required to reduce viability to 50%.

**Supplementary Table 14:** Summary of previous studies of precancers PDO of BO and dysplasia.

| **Study** | **Number of precancerous BO and/or Dysplastic organoid lines** | **Characterisations** |
|---|---|---|
| Long-term expansion of epithelial organoids from human colon, adenoma, adenocarcinoma, and Barrett's epithelium.<br>*Gastroenterology*. 2011 | Two BO lines.<br>Dysplasia not specified | H&E staining with PAS to characterize goblet-like cells |
| Towards personalized induction therapy for esophageal adenocarcinoma: organoids derived from endoscopic biopsy recapitulate the pre-treatment tumor. *Sci Rep,* 2020 | One BO line.<br>Dysplasia not specified<br>Note, this paper primary focused on OAC. Discovery of this Barrett's line is not intentional, but due to the organoids established from a tumour tissue did not harbour TP53 as found in the tissue, therefore deduced as BO line. This study in fact highlighted lack of established framework to identify precancerous organoid lines. | H&E and Alcian Blue staining.<br>IHC: p53 and CK7<br>Genomics: Whole Exome Sequencing (WES) to identify SNVs and Copy Number Variations (CNV). |
| A human Barrett's esophagus organoid system reveals epithelial-mesenchymal plasticity induced by acid and bile salts. *Am J Physiol Gastrointest Liver Physiol*. 2022 | Seven BO lines<br>Dysplasia not specified | H&E and Alcian Blue and PAS staining<br>qRT-PCR on CDX2, KRT7, KERT20, MUC2, SOX9 and LGR5 |
| Molecular characterization of Barrett's esophagus at single-cell resolution, *Proc. Natl. Acad. Sci*. 2021 | Seven BO lines from four patients<br>Two non-dysplastic<br>Five dysplastic | scDNA-seq of organoids, establishing chromosomally stable (CS) and chromosomal instability (CIN) cells. |
| Barrett's Esophagus Epithelial Stem Cells Have Distinct Gene Signatures, *Gastro Hep Advances*, 2024 | 13 organoid lines including: 3 NDBE; 5 LGD; 3 HGD; 2 HGD and EAC | RNA-seq of bulk organoids<br>qRT-PCR on UCA1, FZD10, ST6GAL1, HOXC10, SOX14, CT83, IL23A, MUC7, PTGS2 |

< Supplementary_Tables_6_7_8_15.xlsx>

**Supplementary Table 15:** Metadata relating to archived data for models generated in this study.

